# A fungal endophyte induces local cell-wall mediated resistance in wheat roots against take-all disease

**DOI:** 10.1101/2023.11.23.568424

**Authors:** Tania Chancellor, Daniel P. Smith, Wanxin Chen, Suzanne J. Clark, Eudri Venter, Kirstie Halsey, Esther Carrera, Vanessa McMillan, Gail Canning, Victoria Armer, Kim E. Hammond-Kosack, Javier Palma-Guerrero

## Abstract

Take-all disease, caused by the ascomycete fungus *Gaeumannomyces tritici*, is one of the most important root diseases of wheat worldwide. The fungus invades the roots and destroys the vascular tissue, hindering the uptake of water and nutrients. Closely related non-pathogenic species in the *Magnaporthaceae* family, such as *Gaeumannomyces hyphopodioides*, occur naturally in arable and grassland soils and have previously been reported to reduce take-all disease in field studies. However, the mechanism of take-all protection has remained unknown. Here, we characterise the root infection biologies of *G. tritici* and *G. hyphopodioides* in wheat. We investigate the ultrastructure of previously described “subepidermal vesicles” (SEVs), produced in wheat roots by non-pathogenic *G. hyphopodioides,* but not by pathogenic *G. tritici.* We show that *G. hyphopodioides* SEVs share key characteristics of fungal resting structures; containing a greater number of putative lipid bodies and a significantly thickened cell wall compared to infection hyphae. We demonstrate that take-all control is achieved via local but not systemic host changes in response to prior *G. hyphopodioides* root colonisation. A time-course wheat RNA sequencing analysis revealed extensive transcriptional reprogramming in *G. hyphopodioides* colonised tissues, characterised by a striking downregulation of key cell-wall related genes, including cellulose synthase (CESA), and xyloglucan endotransglucosylase/hydrolase (XTH) genes. In the absence of take-all resistant wheat cultivars or non-virulent *G. tritici* strains, studying closely related non-pathogenic *G. hyphopodioides* provides a much-needed avenue to elucidate take-all resistance mechanisms in wheat.

## Introduction

Wheat (*Triticum aestivum*) is one of the most important cereal crops worldwide, providing around 20% of human caloric intake globally. Sustaining excellent root health is critical for the acquisition of water and essential nutrients. As global temperatures continue to rise, root health is predicted to face increasing threats from various soil-borne fungal pathogens (Delgado-Baquerizo et al., 2020). The necrotrophic fungal pathogen *Gaeumannomyces tritici*, belonging to the *Magnaporthaceae* family, is responsible for take-all disease, one of the most important root problems of wheat crops worldwide (Freeman & Ward, 2004; Palma-Guerrero et al., 2021). The disease drastically diminishes grain yields during heavy infection episodes. However, due to the genetic intractability of *G. tritici*, both the pathogen and the cereal-pathosystem remain understudied by the molecular plant-microbe interaction community. Root-confined vascular infection by *G. tritici* results in the development of characteristic necrotic lesions originating from the stele which severely disrupt root functions, causing premature crop ripening and reduced grain yield/quality (Asher & Shipton, 1981; Huang et al., 2001). Take-all fungal inoculum builds up in the soil following consecutive wheat crops, and though recent surveys of take-all disease levels are lacking, yield losses of up to 60% have been reported in the UK (McMillan et al., 2011). At present, take-all resistant wheat cultivars are not commercially available, and current fungicide seed treatments do not provide complete protection (Freeman et al., 2005).

Understanding root immunity is essential for the development of take-all resistant cultivars. However, the classical model of immunity, characterised by the concerted effect of pathogen-associated molecular pattern (PAMP) triggered immune responses (PTI) and effector triggered immune responses (ETI), is predominantly based on foliar pathogens (Boller & Felix, 2009; Jones & Dangl, 2006; Pok et al., 2022). Roots must constantly interact with a diverse soil microbiome and distinguish pathogenic microbes from, sometimes closely related, non-pathogenic endophytes or beneficial symbionts (Thoms et al., 2021). How plants engage with beneficial microorganisms while restricting damaging pathogens is regarded as one of the top 10 unanswered questions by the molecular plant microbe-interaction (MPMI) research community (Harris et al., 2020). The selective response of plants to microbes with different lifestyles can be partly explained by the compartmentalisation of localised immune responses in roots (Zhou et al., 2020), and the recognition of microbe-associated molecular patterns (MAMPs), damage-associated molecular patterns (DAMPs) and pathogen-associated molecular patterns (PAMPs) by multiple receptors (Thoms et al., 2021). However, further comparative studies into endophytic and pathogenic plant infecting microbes are sorely needed.

Several members of the *Magnaporthaceae* family are classified within the *Gaeumannomyces-Phialophora* complex (Hernández-Restrepo et al., 2016). *Phialophora* species, such as *Gaeumannomyces hyphopodioides,* occur naturally in grasslands and arable field sites, though do not cause disease symptoms in arable crops (Deacon, 1973; Ulrich et al., 2000; Ward & Bateman, 1999). For this reason, such species have been described as “non-pathogenic”. Wheat colonisation by non-pathogenic *Magnaporthaceae* species can be easily distinguished from wheat infection by pathogenic *G. tritici* due to the production of dark swollen fungal cells in the root cortex. The swollen cells measure between 12 µm and 30 µm in diameter, depending on the fungal species (Deacon, 1976a). These enigmatic structures have been previously described as pigmented cells (Holden, 1976), growth cessation structures (Deacon, 1976a) or subepidermal vesicles (SEVs) (Osborne et al., 2018). The closely related rice leaf blast pathogen *Magnaporthe oryzae,* is also reported to form SEV-like structures in cereal roots. *M. oryzae* can infect rice root tissues (Dufresne & Osbourn, 2001; Marcel et al., 2010), producing brown spherical structures resembling SEVs in epidermal and cortical cells. SEVs may form following growth cessation of a hyphal apex (Deacon,1976a).

Prior colonisation by certain non-pathogenic *Magnaporthaceae* species is reported to provide protection against take-all disease in field studies (Wong et al., 1996). Furthermore, Osborne et al., (2018) demonstrated that certain elite winter wheat varieties have an improved ability to promote *G. hyphopodioides* populations in field soils, suggesting that careful cultivar choice during wheat rotations could provide a natural level of biocontrol. However, as far as we are aware, disease protection by non-pathogenic *Magnaporthaceae* species has not been reported in any recent publications, and the mechanism(s) underlying disease control remain unknown. The existence of endophytic species conferring resistance against closely related pathogens is not limited to the *Magnaporthaceae* family. Non-pathogenic strains in the *Fusarium oxysporum* species complex are known to provide protection against Fusarium wilt disease, a major disease caused by pathogenic *F. oxysporum* strains (de Lamo & Takken, 2020). This phenomenon, termed endophyte-mediated resistance (EMR), is reportedly independent of jasmonic acid (JA), ethylene (ET) and salicylic acid (SA) signalling (Constantin et al., 2019).

Here, we provide the first comparative analysis of wheat transcriptional responses to *G. tritici* and *G. hyphopodioides* across key stages of early fungal infection. We characterise the different fungal structures produced and some of the wheat cell wall changes occuring during root infection. Our findings shed light on the distinct plant responses to these two closely related root infecting fungi with contrasting lifestyles, and help to pinpoint localised mechanisms for the control of take-all disease by *G. hyphopodioides*. Together, our findings contribute to an improved understanding of wheat root resistance against take-all disease.

## Results

### Hyphal interactions between *G. hyphopodioides* and *G. tritici*

To investigate the role of direct hyphal interaction in take-all control, a series of fungal confrontation assays were conducted on potato dextrose agar (PDA) plates. Prior to hyphal contact, the individual growth rates of *G. tritici* and *G. hyphopodioides* colonies did not significantly differ from the dual colony controls (table S1). The same was true when the two species were grown in a “sandwich” plate set-up (figure 1A, table S2), suggesting that prior to hyphal contact, neither species produce diffusible antifungal compounds or volatile organic compounds *in vitro*. When hyphae of the two fungal species interacted in confrontation assays, a dark barrage was observed in the interaction zone (figure 1A). The observed barrage formed 1-2 days following hyphal interaction, perhaps suggesting that direct interaction causes hyphal stress in at least one of the interacting species. A dark barrage was not observed when isolates of the same species were confronted (figure S1).

**Figure 1.**
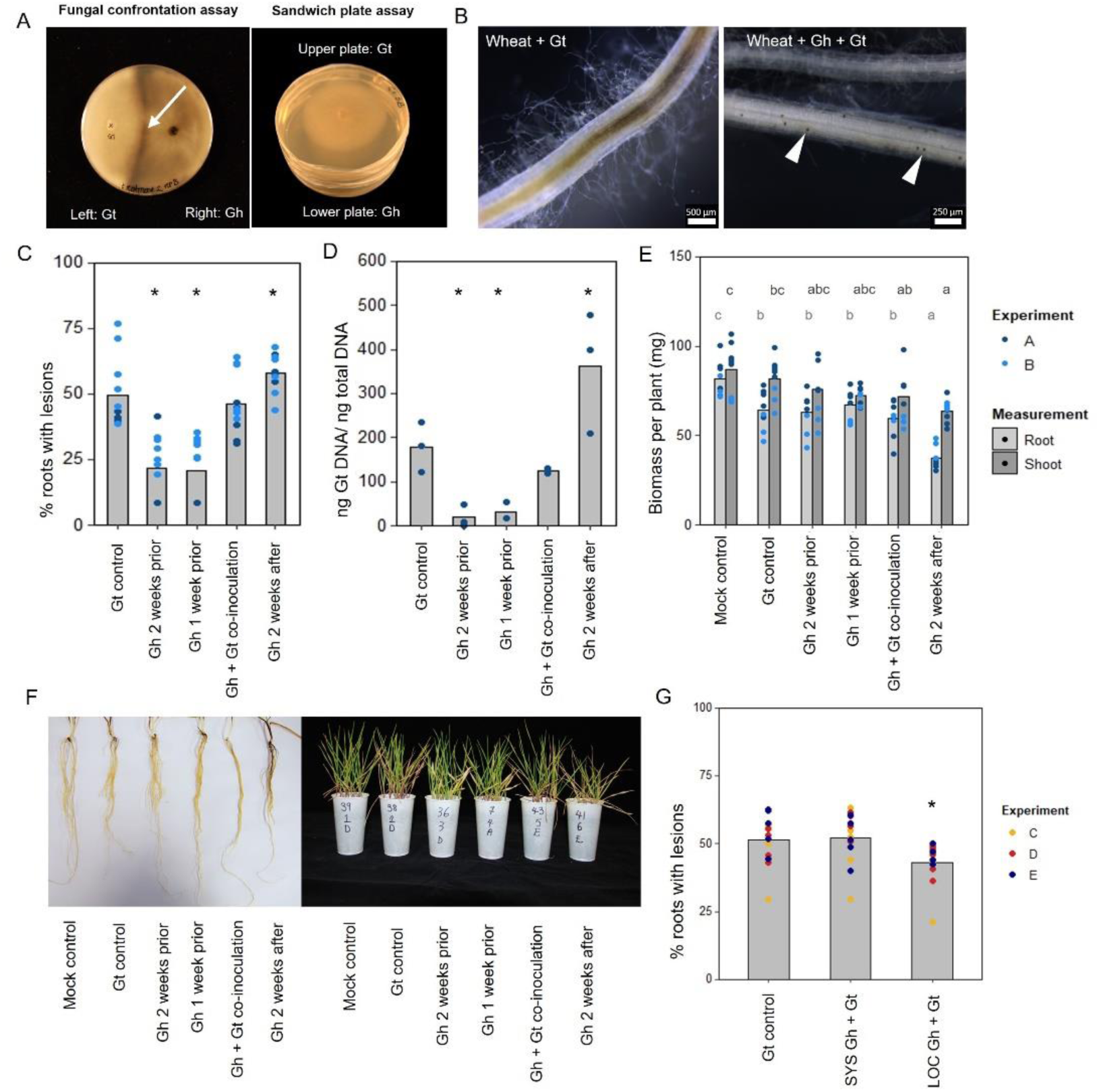
*In vitro* and *in planta* interaction studies involving endophytic *G. hyphopodioides* and pathogenic *G. tritici*. A. *In vitro* fungal interaction assays on PDA plates. Fungal confrontation assays imaged 2 days following hyphal interaction, colonies in sandwich plate assays imaged 6 days after establishment. Arrow indicates a dark barrage in the interaction zone; B. Stereomicroscope images of wheat roots infected with *G. tritici* only, or co-inoculated with *G. hyphopodioides.* Arrowheads indicate *G. hyphopodioides* sub-epidermal vesicles (SEVs); C. Percentage of wheat roots (cv. Hereward) with take-all root lesions in co-inoculation experiments with *G. hyphopodioides* (GLM: F=25.99, d.f. 4, 49, p <0.001); D. *G. tritici* fungal biomass (ng *G. tritici* DNA/ ng total DNA) in co-inoculation experiments with *G. hyphopodioides,* as quantified by qPCR (F=61.10, d.f. 4, 38, p<0.001); E. Shoot and root dry biomass (mg) in *G. tritici* co-inoculation experiments with *G. hyphopodioides* (F=6.49, d.f. 5, 36, p<0.001; F=4.50, d.f. 5, 36, p<0.01, respectively); F. Representative images of wheat roots (left) and shoots (right) in co-inoculation experiments; G. The percentage of wheat roots (cv. Chinese Spring) with take-all lesions in split root co-inoculation experiments with *G. hyphopodioides* (F=29.44, d.f. 2, 36, p=0.007). Asterisks indicate a significant difference to the *G. tritici* control as calculated by Dunnett’s post-hoc test (p<0.05). Letters indicate significant differences as calculated by Tukey’s multiple comparisons test (p<0.05). Gt= *G. tritici,* Gh= *G. hyphopodioides*, SYS=systemic, LOC=local.

### Pre-treatment with *G. hyphopodioides* provides local control against take-all disease

To investigate the hypothesis that non-pathogenic *G. hyphopodioides* provides protection against take-all disease by inducing wheat resistance, seedling co-inoculation experiments were carried out under controlled environment conditions. *G. hyphopodioides* inoculum was added to wheat seedlings (cv. Hereward) growing in pots 1 week prior, 2 weeks prior, at the same time as, and 1 week after inoculation with pathogenic *G. tritici.* Characteristic black necrotic root lesions were observed in *G. tritici* infected control plants. SEVs were observed in plants co-inoculated with *G. hyphopodioides* (figure 1B). The data revealed a significant reduction in both take-all disease levels and *G. tritici* fungal biomass in plants pre-treated with *G. hyphopodioides* 2-weeks prior or 1-week prior to *G. tritici* inoculation (figure 1C, D). Hence, even very early colonisation by *G. hyphopodioides* is sufficient for take-all control. These findings were consistent with additional experiments involving a different *G. tritici* isolate (Gt 17LH(4)19d1) and wheat cultivar (cv. Chinese Spring) (figure S2).

Seedlings co-inoculated with *G. hyphopodioides* and *G. tritici* at the same time had no effect on take-all disease levels or *G. tritici* fungal biomass. Importantly, adding *G. hyphopodioides* after *G. tritici* resulted in increased levels of take-all disease and *G. tritici* fungal biomass (figure 1C, D). Furthermore, the shoot and root dry biomass of plants in this latter treatment were significantly reduced, indicating that seedling health is negatively affected when *G. hyphopodioides* infections occur in addition to *G. tritici* infection (figure 1E, F). These findings should be taken into careful consideration when evaluating the potential of *G. hyphopodioides* as a biocontrol agent.

Split-root experiments were carried out to determine whether *G. hyphopodioides* provides local or systemic protection against take-all disease. Significant disease reduction was achieved only in roots which had been directly inoculated with *G. hyphopodioides* (LOC), and not in systemic roots (SYS) which had not been directly inoculated with *G. hyphopodioides* (figure 1G). Taken together, we demonstrate that local induced wheat resistance plays a crucial role in the control of take-all disease by *G. hyphopodioides,* and this response is consistent across both winter and spring wheat types.

### The differing infection biologies *of G. hyphopodioides* and *G. tritici* in wheat roots

To study fungal infection processes during early root colonisation, wheat seedlings (cv. Chinese Spring) were root inoculated with either *G. hyphopodioides* (NZ.129.2C.17) or *G. tritici* (Gt 17LH(4)19d1) in an agar plate system. Plants were harvested at 2, 4 and 5 dpi to capture key stages of fungal infection for later RNA-seq analysis. At 2 dpi, very few hyaline runner hyphae were detected on the root surface of plants inoculated with either *G. tritici* or *G. hyphopodioides*, and hyphae had not yet penetrated the epidermal cells in either interaction (figure 2A, B). By 4 dpi, hyaline runner hyphae covered a greater area of the root surface (figure S3) and hyphae were detected in epidermal and cortical cells of both *G. tritici* and *G. hyphopodioides* inoculated roots (figure 2A, B). At 5 dpi, hyaline runner hyphae were detected across a large area of the root surface (figure S3) in both fungal treatments. *G. tritici* hyphae infected the stele, whereas *G. hyphopodioides* hyphal growth was arrested in the cortex (figure 2B). *G. hyphopodioides* hyphae were detected in cortical cells, from which SEVs were formed (figure 2A). Newly formed SEVs could be visualised by wheat germ agglutinin (WGA) staining, whereas mature SEVs, which were darker in colour, could not be visualised by WGA staining (figure S4). *G. tritici* did not produce SEVs in wheat roots at any time point, and *G. hyphopodioides* hyphae were not observed in the stele at any time point (figure S5).

**Figure 2.**
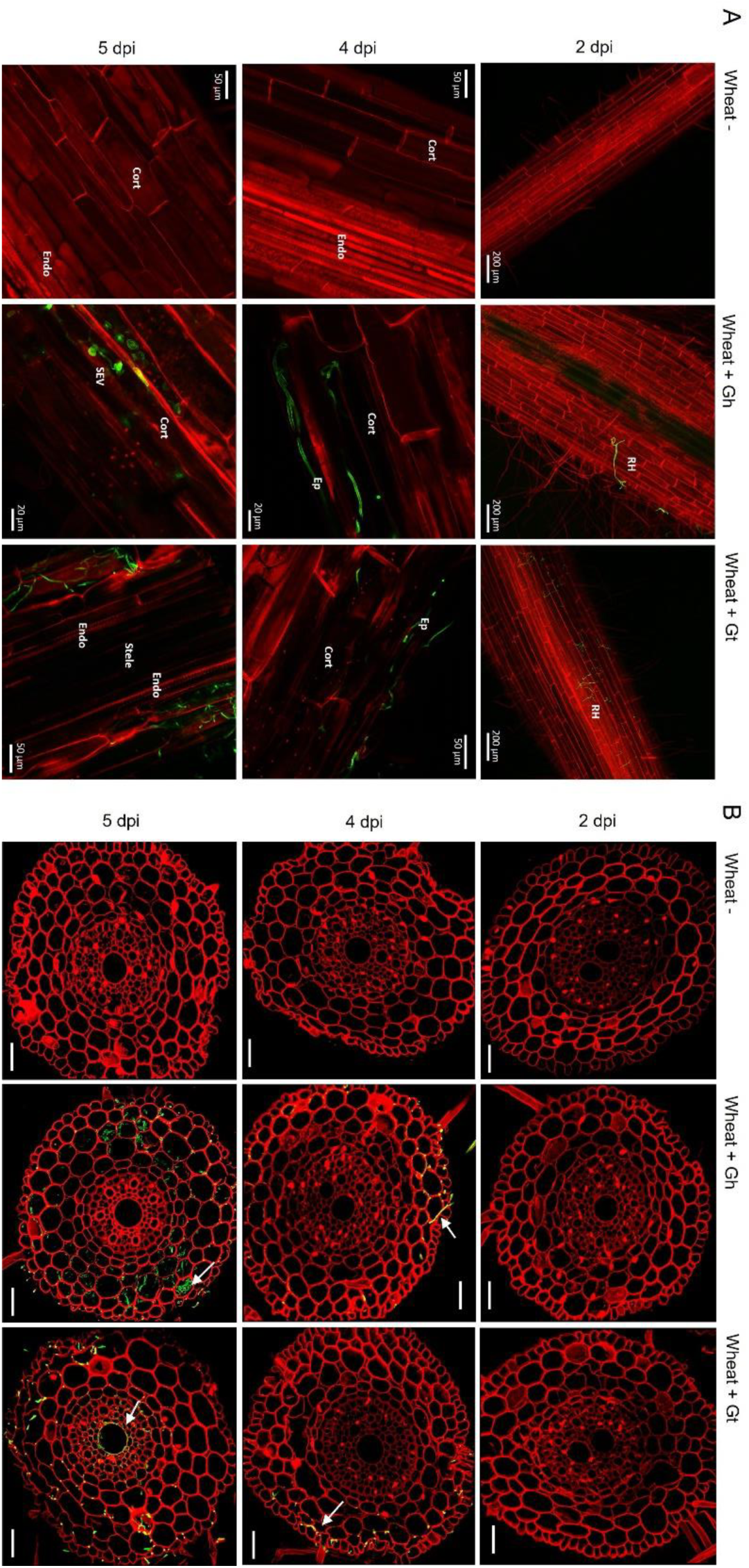
Fluorescence images obtained by confocal microscopy of mock inoculated*, G. hyphopodioides* colonised or *G. tritici* infected wheat roots. A. Confocal micrographs of whole root pieces highlighting fungal infection structures; B. Z-stack images of transversal sections showing colonisation of different root cell layers across time points. Gt= *G. tritici,* Gh=*G. hyphopodioides,* RH= runner hyphae, Ep= epidermal cell, Cort= cortical cell, SEV= subepidermal vesicle, Endo= endodermal barrier. Fungal hyphae (green) are stained with WGA-AF488, plant cell-walls (red) are stained with propidium iodide. White arrows in panel B indicate fungal hyphae. Scale bars in panel B represent 50 µm.

To investigate the structure of mature SEVs, wheat plants (cv. Hereward) were inoculated with *G. hyphopodioides* (NZ.129.2.17) in a seedling pot infection assay. Colonised plants were harvested at 5 weeks post inoculation and imaged by transmission electron microscopy (TEM). Comparative analysis of intraradical fungal hyphae and SEVs revealed that SEVs contain a greater number of putative lipid bodies and a significantly thickened cell wall, comprising two to three layers of differing densities (figure 3A, B). Multiple SEVs were often observed in a single plant cell (figure 3C) and SEVs were often found appressed to the plant cell wall (figure 3 D, E).

**Figure 3.**
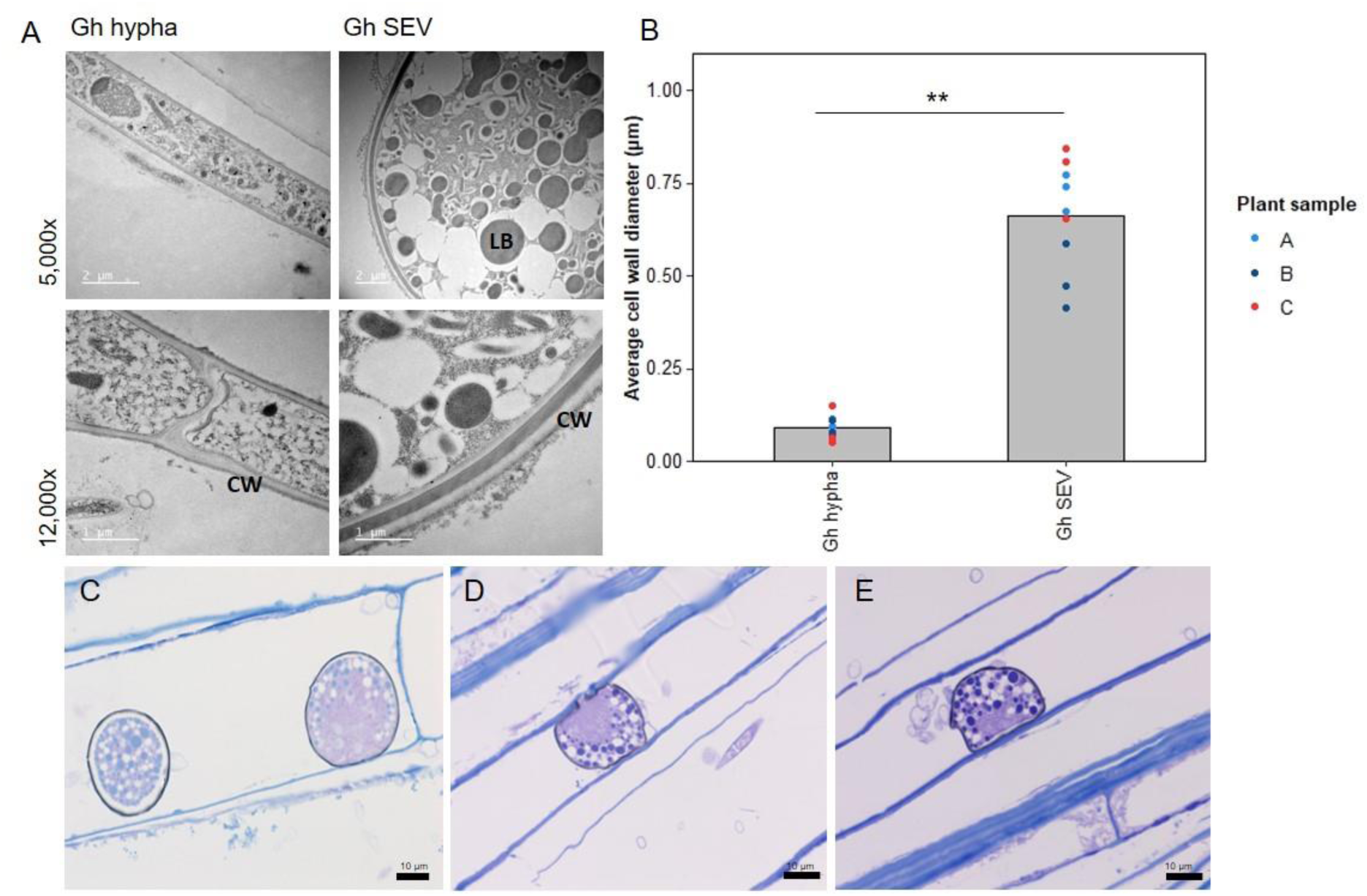
Subepidermal vesicles (SEVs) produced by *G. hyphopodioides* (Gh) in wheat roots. A. Transmission electron micrographs (TEM) of *Gh* hyphae (left) and SEVs (right); B. Average cell wall diameter (µm) of *Gh* fungal structures (F= 8.3, d.f. 2, 12, p<0.01); C-E. Light micrographs of *Gh* SEVs in semi-thin sections, stained with toluidine blue. CW=plant cell wall; LB=putative lipid body.

### Wheat transcriptional remodelling during fungal infection

Three time-points (2, 4 and 5 dpi) were selected for RNA-seq analysis based on the stage of fungal infection (figure S5). Principal Component Analysis (PCA) of sample distances demonstrated a good level of clustering between biological replicates, though *G. tritici* infected samples exhibited comparatively higher levels of variation (figure 4A). Gene expression levels were compared between *G. tritici* infected or *G. hyphopodioides* colonised plants and the uninoculated control plants at each time point individually. Full lists of the differentially expressed genes (DEGs) can be found in table S3. As expected, the number of wheat DEGs between the uninoculated control and *G. tritici* infected or *G. hyphopodioides* colonised plants was low at 2 dpi (77 and 62, respectively). By 4 dpi, *G. tritici* infection and *G. hyphopodioides* colonisation resulted in the differential expression of 1061 and 1635 wheat genes, respectively. At 5 dpi, a striking number of wheat genes were DE in response to *G. hyphopodioides* colonisation (7532), whereas the number of DEGs in response to *G. tritici* infection (1074) showed little change compared to 4 dpi (figure 4B).

**Figure 4.**
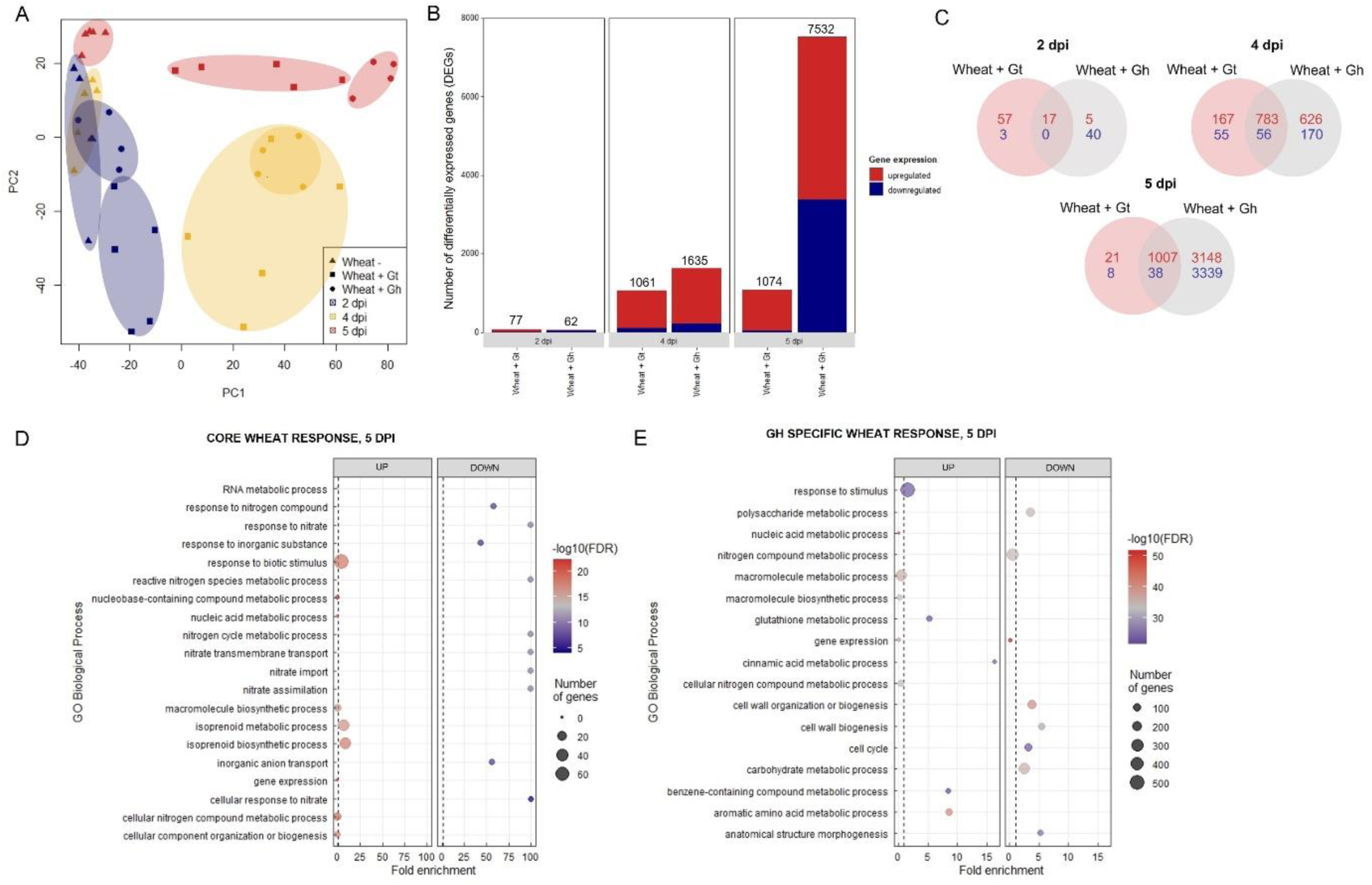
Transcriptional profiling of *G. hyphopodioides* colonised or *G. tritici* infected wheat roots. A. Principal Component Analysis (PCA) plot of sample distances based on transformed (variance stabilising transformation) gene count data. Data points have been categorised by shape and colour to denote treatment and time point, respectively; B. The number of differentially expressed genes (DEGs) in wheat colonised by *G. hyphopodioides* or *G. tritici* compared to uninoculated control samples; C. Venn diagram highlighting the number of unique and shared wheat DEGs in *G. tritici* infected or *G. hyphopodioides* colonised samples compared to the uninoculated control samples; D. Top 10 enriched biological process GO terms among DEGs in the shared wheat response to both *G. tritici* and *G. hyphopodioides* at 5 dpi; E. Top 10 enriched biological process GO terms unique to the wheat response to *G. hyphopodioides* colonisation at 5 dpi. The top 10 GO terms were determined by false discovery rate (FDR).

To investigate wheat transcriptional changes during the infection progression of *G. hyphopodioides* compared to *G. tritici*, gene ontology (GO) enrichment analyses were carried out on the sets of DEGs described above. At 2 dpi, genes involved in the terpenoid biosynthetic process/metabolic process were upregulated in *G. tritici* inoculated roots. Meanwhile, genes involved in the nicotianamine biosynthetic process/metabolic process were downregulated in *G. hyphopodioides* inoculated roots (figure S6A, B). At 4 dpi, genes involved in the cinnamic acid biosynthetic/metabolic process and the L-phenylalinine metabolic/catabolic process were upregulated in *G. hyphopodioides* colonised wheat roots, suggesting that lignin biosynthesis is important at this time point. However, these GO terms were not significantly enriched until 5 dpi in *G. tritici* infected roots, suggesting that lignin biosynthesis is also involved in the defence response to *G. tritici*, though at a later stage than *G. hyphopodioides.* Other enriched terms in *G. hyphopodioides* colonised plants at 5 dpi included response to wounding, regulation of defence responses and regulation of the jasmonic acid (JA) signalling pathway. Downregulated terms included gene expression, plant-type cell wall organisation or biogenesis and RNA metabolic process (figure S6A, B).

Next, we compared the unique and shared wheat transcriptional responses to the two fungal species. At 5 dpi, 97% of the genes which were DE in response to *G. tritici* infection were also DE in response to *G. hyphopodioides* colonisation (figure 4C). Within this core set of genes at 5 dpi, highly enriched GO biological process terms included isoprenoid biosynthesis, plant response to biotic stimulus and isoprenoid biosynthetic/metabolic process (figure 4D). The plant response to biotic stimulus term comprised 42 DE genes, 14 of which encoded proteins containing small cysteine-rich protein (SCP)-domains, often associated with pathogenesis-related proteins. Six genes encoded chitinases, two encoded wound induced proteins (WIN) and a further four encoded protein kinase domain-containing proteins, thus indicating a clear defence response to both fungi (table S4). Enriched biological process GO terms among shared downregulated genes included response to nitrate, nitrate transmembrane transport and nitrate assimilation (figure 4D). Highly enriched molecular function GO terms among upregulated genes included manganese ion binding, oxidoreductase activity and heme binding. Highly enriched molecular function GO terms among downregulated genes included nitrate transmembrane transporter activity and oxygen binding (table S5A). Highly enriched cellular component GO terms among upregulated genes included extracellular region (table S5B).

GO enrichment analysis was repeated for DEGs unique to the wheat response to *G. hyphopodioides* at 5 dpi. Cinnamic acid metabolic process, glutathione metabolic process and benzene-containing compound metabolic process were among the top 10 upregulated biological process GO terms. In contrast, cell wall organisation or biogenesis, cell cycle and anatomical structure morphogenesis were among the top 10 downregulated biological process GO terms (figure 4E). Highly enriched molecular function GO terms among upregulated genes included phenylalanine ammonia lyase (PAL) activity, glutathione transferase activity and ion binding. In contrast, structural constituents of chromatin, tubulin binding and nucleosome binding were highly enriched among the downregulated genes (table S6A). Highly enriched cellular component function GO terms among downregulated genes included nucleosome, microtubule cytoskeleton and protein-DNA complex (table S6B).

### Wheat phytohormone response to *G. hyphopodioides* colonisation and *G. tritici* infection

Regulation of the JA signalling pathway was identified as a newly upregulated GO term at 5 dpi in *G. hyphopodioides* colonised roots (see above). The GO term comprised 26 DEGs (out of a total of 77 known genes in wheat), all of which were TIFY transcription factors. In contrast, just three TIFY transcription factors (TIFY10C-like_TraesCS5D02G219300, TIFY10C-like_TraesCS5B02G211000, TIFY11E-like_ TraesCS7D02G204700) were DE in *G. tritici* infected plants compared to the control. Although not identified by GO enrichment analysis, we also investigated the expression of JA biosynthesis genes. In total, 23 JA biosynthesis-related genes were DE (17 up/ 6 down) in response to *G. hyphopodioides* at 5 dpi. The list included lipoxygenase (LOX), allene oxide synthase (AOS) and AOS-like, 12-oxophytodienoate reductase (OPR) and OPR-like, and 3-ketoacyl-CoA thiolase (KAT-like) genes. Of these genes, only one (LOX8_TraesCS7B02G145200) was differentially expressed in response to *G. tritici* at 5 dpi (figure 5A, table S7).

**Figure 5.**
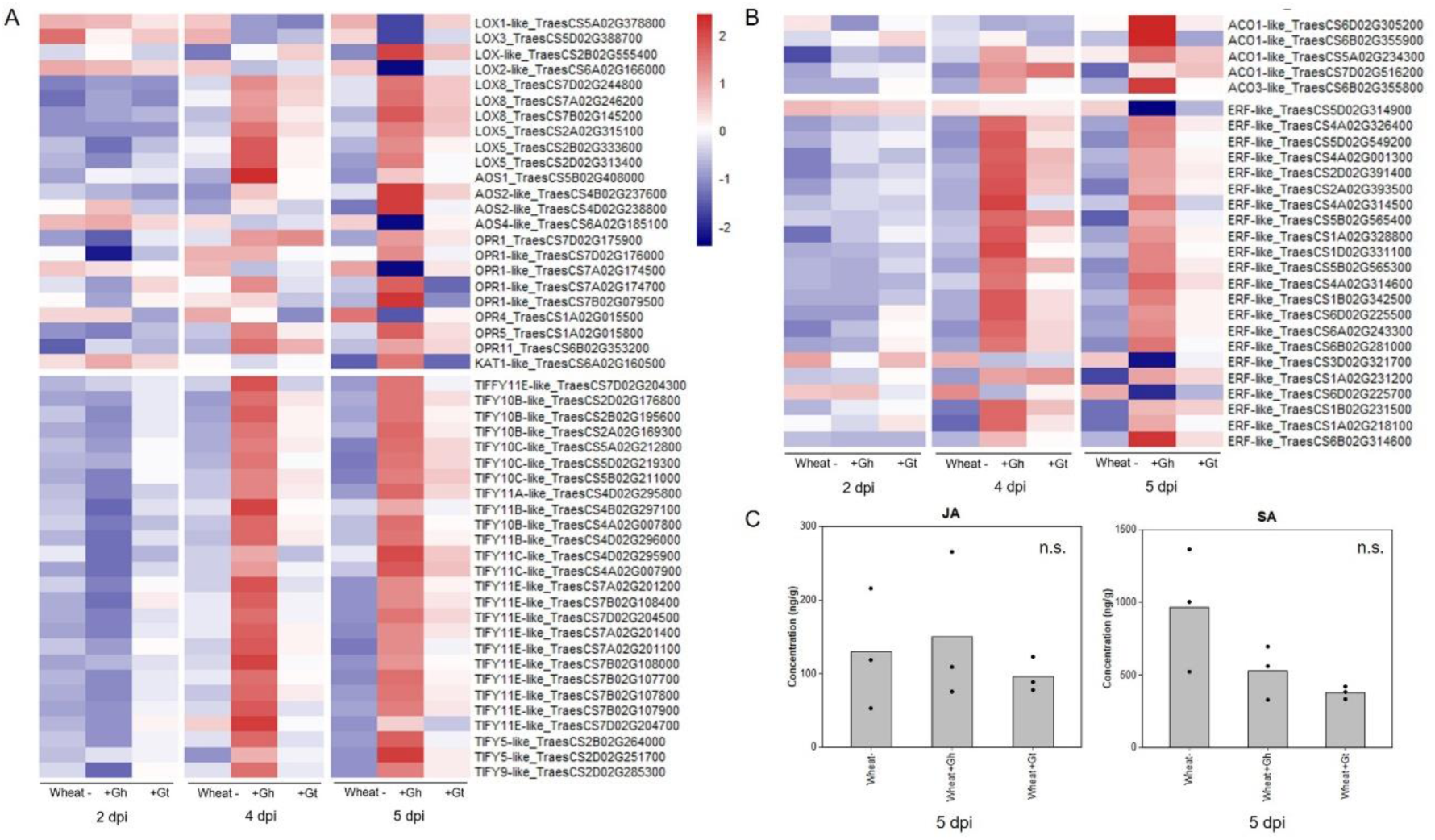
Wheat phytohormone-associated gene responses and JA quantification in response to *G. hyphopodioides* (Gh) colonisation or *G. tritici* (Gt) infection. A. Expression of genes involved in the biosynthesis of JA and the regulation of the JA mediated signalling pathway; B. Expression of genes involved in ET biosynthesis and downstream ET signalling pathways. Heatmap data represent LOG transformed normalised genes counts; C. Phytohormone quantification of JA and SA in roots harvested at 5 dpi (F=0.30, d.f. 2, 6, p= 0.75; F=4.19, d.f. 2, 6, p=0.07, respectively). n.s.= not significant. Data have been back-transformed from a square root scale. ACO, 1-aminocyclopropane-1-carboxylic acid oxidase; AOC, allene oxide cyclase; AOS, allene oxide synthase; ERF, ethylene responsive transcription factor; KAT, 3-ketoacyl-CoA thiolase; LOX, lipoxygenase; OPR, 12-oxophytodienoate reductase; TIFY, TIFY-domain containing transcription factor.

The JA and ET signalling are often closely linked. Therefore, we investigated genes involved in ET biosynthesis and signalling. Five ACC-oxidase (ACO-like) genes were upregulated in response to *G. hyphopodioides* at 5 dpi. In addition, 22 ethylene responsive transcription factor-like (ERF-like) genes, key integrators of downstream ET and JA signal transduction pathways (Lorenzo et al., 2003), were DE (19 up/ 3 down) in response to *G. hyphopodioides* by 5 dpi. In contrast, four ERF-like genes (TraesCS4A02G001300, TraesCS5B02G565400, TraesCS1A02G231200, TraesCS1B02G231500) were upregulated at 4 dpi and two (TraesCS5B02G565400, TraesCS1A02G231200) were upregulated at 5 dpi in *G. tritici* infected roots compared to the control (figure 5B, table S7). Salicylic acid (SA) is another key phytohormone involved in the plant response to pathogen invasion. SA signalling was not identified as a significantly enriched GO term in response to *G. hyphopodioides* or *G. tritici* at any time point.

To investigate whether the local transcriptional changes described above resulted in altered hormone levels, hormone quantifications of JA and SA were carried out in *G. hyphopodioides* colonised, *G. tritici* infected and uninoculated control roots at 5 dpi. We found no significant difference in the levels of JA or SA between any treatments (figure 5C).

### *G. hyphopodioides* colonisation results in the early induction of lignin biosynthesis genes

PAL activity, essential for the lignin biosynthesis pathway, was identified as a significantly enriched molecular function GO term in the unique wheat response to *G. hyphopodioides* at 5 dpi (see table S6A). To investigate root lignification in response to *G. hyphopodioides* and *G. tritici*, we explored the expression of key genes involved in the lignin biosynthesis pathway in wheat (figure 6A). *G. hyphopodioides* colonisation resulted in the earlier upregulation of lignin biosynthesis genes compared to *G. tritici*, with key genes such as arogenate dehydratase (ADT), phenylalanine ammonia-lyase (PAL), cinnamate 4-hydroxylase (4CL) and cinnamoyl-CoAreductase (CCR) significantly upregulated at 4 dpi (figure 6B). However, two of eight caffeic acid *O*-methyltransferase (COMT) genes detected were already significantly upregulated in response to *G. tritici* at 2 dpi (TraesCS5D02G488800, TraesCS5D02G488900). Interestingly, the remaining COMT genes (TraesCS2B02G066100, TraesCSU02G024300, TraesCS3B02G612000, TraesCS7D02G539100, TraesCS6D02G008200, TraesCS7D02G538900) were strongly downregulated in response to *G. hyphopodioides* by 5 dpi, suggesting a decrease in the proportion of syringyl (S)-lignin. Most striking however, was the significant upregulation of 37 PAL genes in response to *G. hyphopodioides*, compared to the upregulation of just 12 PAL genes in response to *G. tritici* at 5 dpi (figure 6B, table S8).

**Figure 6.**
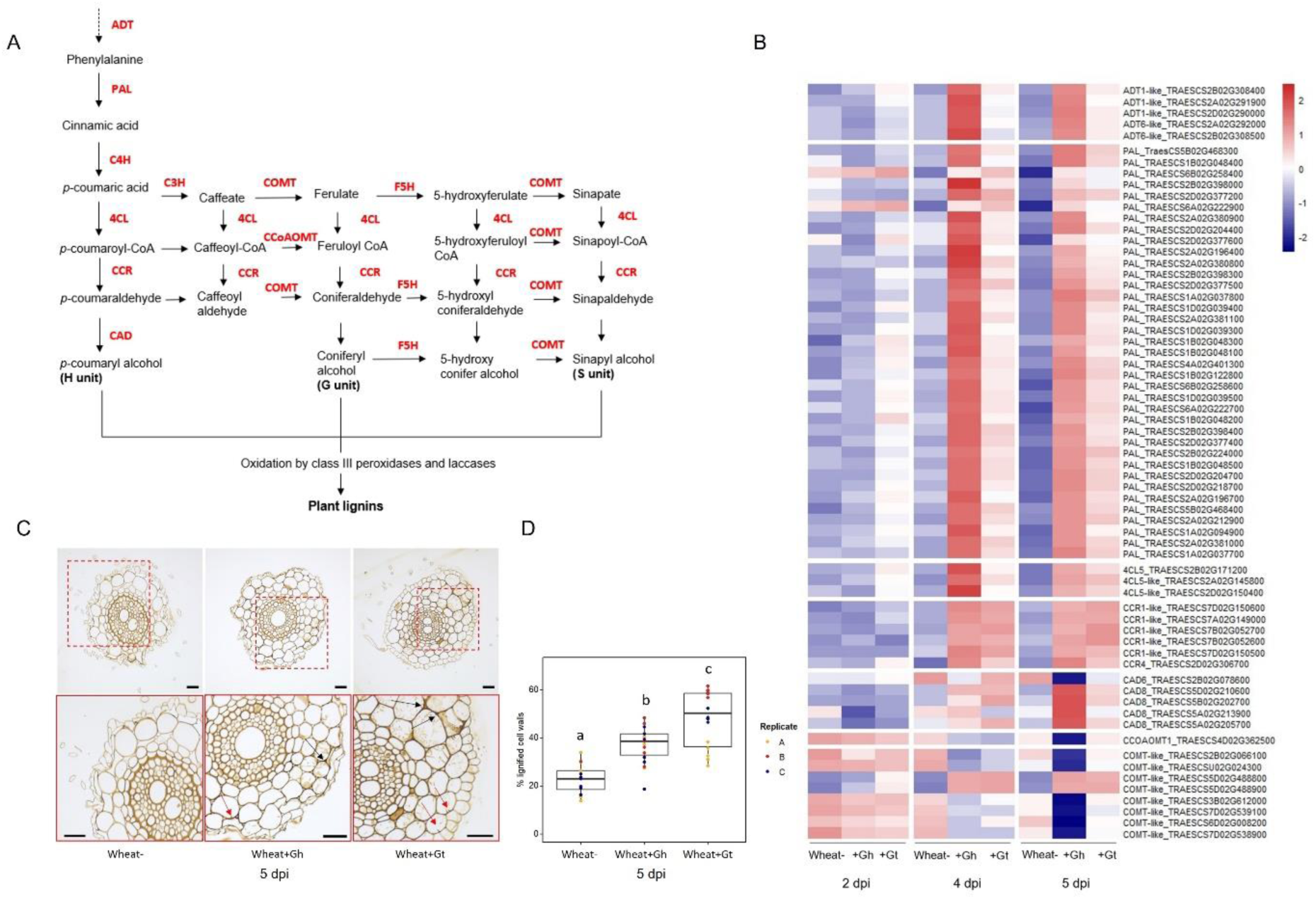
Lignin biosynthesis pathway and the lignin responses to *G. hyphopodioides* colonisation or *G. tritici* infection. A. Schematic of the lignin biosynthesis pathway in plants (adapted from Nguyen et al., 2016); B. Expression of key genes involved in the lignin biosynthesis pathway, based on LOG transformed normalised gene counts; C. Micrographs of transversal root sections stained with potassium permanganate for the visualisation of cell wall lignification in response to fungal infection at 5 dpi. Black arrows indicate lignified cell wall thickenings, red arrows indicate plant lignitubers. Scale bars represent 50 µm; D. Mean percentage of total cell wall area stained within dark parameters, indicating relative cell wall lignification (F=34.61, d.f. 2, 50, p<.001). Lowercase letters indicate Tukey post-hoc groupings. ADT, Arogenate dehydratase; PAL, phenylalanine ammonia-lyase; C4H, cinnamate 4-hydroxylase; 4CL, 4-coumarate CoA ligase; HCT, quinateshikimate *p*-hydroxycinnamoyltransferase; C3’H, *p*-coumaroylshikimate 3′-hydroxylase; CCoAOMT, caffeoyl-CoA *O*-methyltransferase; CCR, cinnamoyl-CoAreductase; F5H, ferulate 5-hydroxylase; CAD, cinnamyl alcohol dehydrogenase; COMT, caffeic acid *O*-methyltransferase.

To visualise lignification of infected root tissues, potassium permanganate staining was performed on transverse sections of samples harvested at 5 dpi (figure 6C). The percentage of total cell wall area with dark potassium permanganate staining (measured in ImageJ) was used to quantify relative cell wall lignification. Based on these measurements, *G. tritici* infected roots exhibited the highest levels of cell wall lignification, though both *G. hyphopodioides* and *G. tritici* infection resulted in increased cell wall lignin levels compared to uninoculated control roots at 5 dpi (figure 6D). Plant lignitubers, lignified callose deposits surrounding hyphal tips (Bradshaw et al., 2020; Huang et al., 2001; Park et al., 2022), were often detected in cells containing fungal hyphae, though these structures were more common in *G. tritici* infected samples.

### *G. hyphopodioides* colonisation results in local downregulation of cell wall organisation and biogenesis genes

The biological process GO term “cell wall organisation and biogenesis”was identified as being unique to the wheat response to *G. hyphopodioides* at 5 dpi (see figure 4E). In total, 124 genes involved in cell wall organisation and biogenesis (out of 1122 total known genes involved in cell wall organisation and biogenesis in wheat) were downregulated in response to *G. hyphopodioides* at 5 dpi (table S9). In contrast, *G. tritici* infection did not lead to the differential expression of any genes within the cell wall organisation and biogenesis GO term. Focussing on the top 30 DE genes within this GO term in *G. hyphopodioides* colonised plants at 5 dpi, we identified six xyloglucan endotransglycosylases/hydrolases (XTH) genes, three cellulose synthase-like A-like (CSLA) genes, one cellulose synthase-like F (CSLF) gene and three fasciclin-like arabinogalactan (FLA) genes (figure 7A). Though just one cellulose synthase-like (CESA) gene was present in the list of top 30 DEGs, a total of 10 CESA genes were downregulated at 5 dpi (table S9). To validate gene expression in the RNA-seq dataset, we identified key cell-wall related genes where all three wheat homoeologues were downregulated in *G. hyphopodioides* colonised plants relative to the mock inoculated controls (figure 7B). RT-qPCR analyses of the selected targets revealed that, as expected, cell-wall related genes CESA7-like, COBL-5D and FLA11 were significantly downregulated in *G. hyphopodioides* colonised plants compared to the mock inoculated controls (figure 7C).

**Figure 7.**
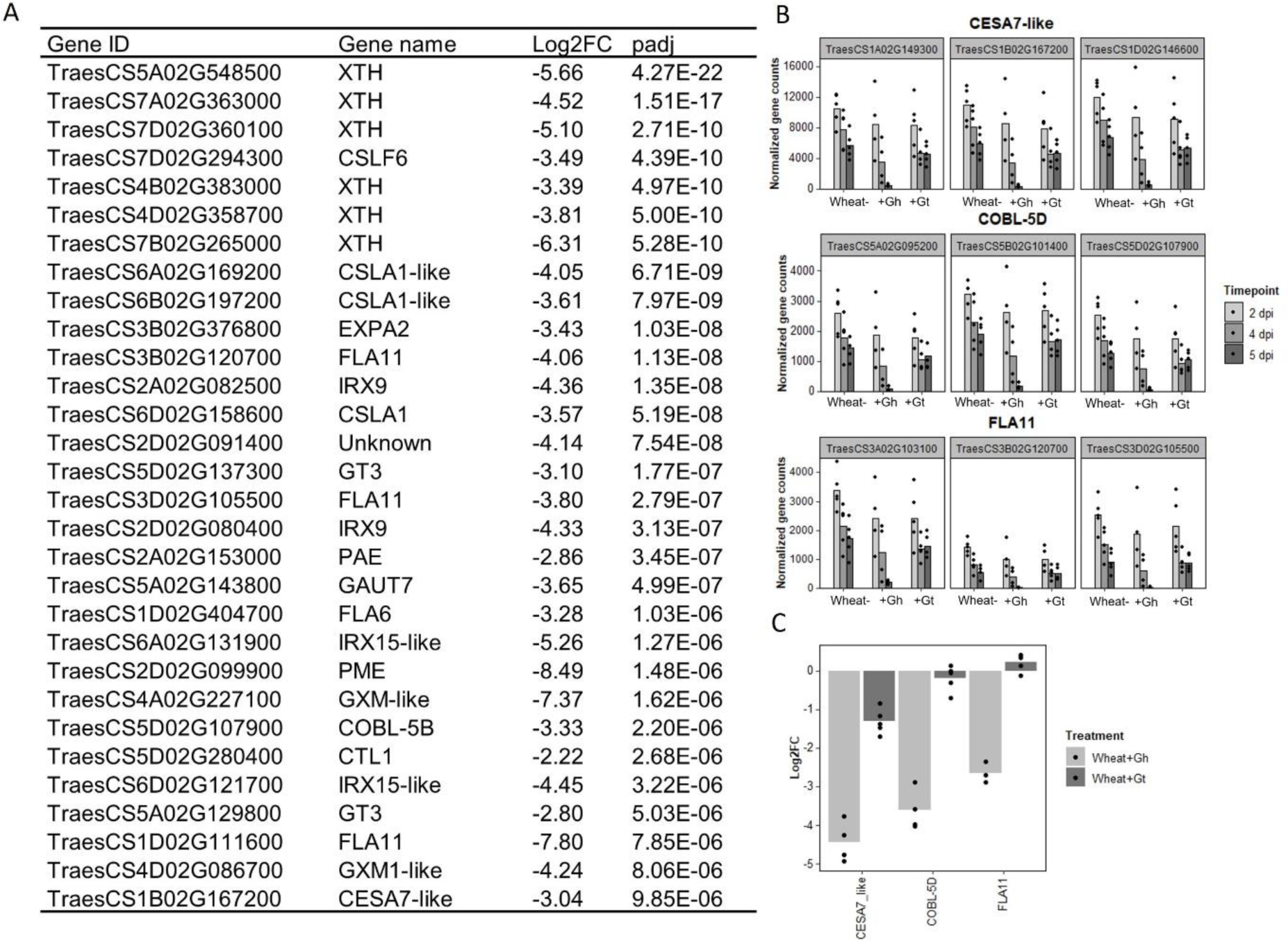
Colonisation by *G. hyphopodioides* results in local downregulation of cell-wall related genes. A. The top 30 most differentially expressed genes relating to cell wall organisation and biogenesis in *G. hyphopodioides* colonised plants at 5 dpi. Ordered by significance (padj); B. Gene counts of selected genes across time points; C. qPCR expression analysis of CESA7-like, COBL-5D and FLA11 in *G. hyphopodioides* colonised or *G. tritici* infected plants at 5 dpi (Log2FC relative to the mock inoculated control).

## Discussion

The biocontrol potential of several non-pathogenic *Magnaportha*ceae species has been reported as early as the 1970s (Deacon, 1973, 1976b; Wong & Southwell, 1980). However, the precise mechanism(s) of control and the molecular pathways underpinning these interactions have remained underexplored. In this study, we show that induced wheat resistance mechanisms play a key role in *G. hyphopodioides*-mediated disease reduction. Furthermore, we demonstrate that these resistance mechanisms operate at a local scale, with effective disease protection conferred in roots pre-treated with *G. hyphopodioides*. However, adding *G. hyphopodioides* after *G. tritici* resulted in increased take-all disease levels, thereby posing a significant risk for field application. The potential for *G. hyphopodioides* to become pathogenic in wheat and/or other cereal crops requires careful investigation. Nevertheless, farmers may exploit the disease suppression ability of *G. hyphopodioides* by growing wheat cultivars known to support natural populations, particularly when placed early in wheat rotations (Osborne et al., 2018).

Though transcriptional studies into the *G. tritici*-wheat interaction have been carried out previously for both host (Kang et al., 2019; Yang et al., 2015; Zhang et al., 2020) and pathogen (Gazengel et al., 2020; Kang., 2019), this is not the case for the *G. hyphopodioides*-wheat interaction. To investigate early wheat responses to *G. tritici* infection and uncover the local wheat defence mechanisms responsible for *G. hyphopodiodes-*induced disease control, we performed comparative transcriptome profiling of *G. hyphopodioides* colonised and *G. tritici* infected wheat using a precision inoculation method. Through detailed screening of infected root material by confocal microscopy, we were able to characterise infection progression across several time-points. In support of early studies into non-pathogenic *Magnaporthaceae* species (Holden, 1976; Speakman & Lewis, 1978), we observed that while pathogenic *G. tritici* grew into the vascular tissues of wheat at 5 dpi, growth of endophytic *G. hyphopodioides* was always limited to the inner cortex. In addition, we observed the formation of *G. hyphopodioides* SEVs in cortical cells at 5 dpi, while SEVs were not observed in *G. tritici* infected roots at any time point. Interestingly, the formation of *G. hyphopodioides* SEVs at 5 dpi was concomitant with a dramatic increase in the number of wheat DEGs. In contrast, the number of wheat DEGs in *G. tritici* infected roots showed minimal increase between 4 dpi and 5 dpi.

TEM analysis of mature *G. hyphopodioides* SEVs revealed that SEVs share key similarities with fungal resting structures such as chlamydospores, both being characterised by a significantly thickened, multi-layered cell wall and an increased number of putative lipid bodies (Francisco et al., 2019). Therefore, we hypothesise that *G. hyphopodioides* SEVs are fungal resting structures which may be produced as a stress response to locally induced host defences, as indicated by extensive transcriptional reprogramming at 5 dpi. Further investigations are required to test this hypothesis and to determine what function, if any, SEVs may play in fungal root infection. In contrast, *G. tritici* infections resulted in far fewer DEGs at 5 dpi (1074), the majority of which were upregulated. Interestingly, almost all DEGs identified in response to *G. tritici* infection were also shared with the wheat response to *G. hyphopodioides* colonisation. Despite triggering a significant wheat defence response, *G. tritici* successfully causes disease, suggesting an ability to either suppress or overcome the local wheat defences triggered. Therefore, future studies should focus on the elucidation of *G. tritici* effectors, enzymes and secondary metabolites, which no doubt contribute to *G. tritici* pathogenicity. One such effector, the ortholog of the barley powdery mildew effector gene BEC1019, has already been associated with *G. tritici* virulence in wheat (Zhang et al., 2019).

Strikingly, 11% of all known cell wall organisation/biogenesis related genes in wheat were downregulated in *G. hyphopodioides* colonised plants at 5 dpi, while none were significantly downregulated in response to *G. tritici.* Impairment of cell wall integrity (CWI) by pathogen invasion triggers the release of antimicrobial compounds and Damage-Associated Molecular Patterns (DAMPs), the latter inducing plant innate immune responses upon recognition by plant Pattern Recognition Receptors (PRRs) (Miedes et al., 2014). In this study, *G. hyphopodioides* colonisation triggered the downregulation of 13 fasciclin-like arabinogalactan (FLA) genes and 18 xyloglucan endotransglucosylase/hydrolase (XTH) genes. FLA proteins contain a putative cell adhesion domain which may link the cell membrane and the cell wall. FLA mutants in Arabidopsis exhibit a range of phenotypes including reduced cellulose content, altered secondary cell-wall deposition and reduced tensile strength (Ashagre et al., 2021). XTH genes are also involved in the maintenance of CWI; these genes encode xyloglucan modifying enzymes which cleave xyloglucan chains to enable cell wall expansion or alter cell wall strength (Cosgrove, 2005). In addition, we detected the downregulation of 10 CESA genes. Though the exact mechanism is not yet known, a number of studies in Arabidopsis indicate a link between CESA expression, CWI and disease resistance. Mutations in the CESA4, CESA7 and CESA8 genes, required for secondary cell wall formation in Arabidopsis, confer enhanced resistance to the necrotrophic fungus *Plectrosphaerella cucumerina* and the biotrophic bacterium *Ralstonia solanacearum* (Hernández-Blanco et al., 2007). In addition, pathogenic *Fusarium oxysporum* root infection of Arabidopsis results in the downregulation of various CESAs, causing an alteration in primary cell wall cellulose and contributing to disease resistance (Menna et al., 2021). Furthermore, mutations in CESA genes in Arabidopsis trigger the activation of defence responses and the biosynthesis of lignin, regulated at least in part, by the jasmonic acid (JA) and ethylene (ET) signalling pathways (Caño-Delgado et al., 2003). A link between JA/ET signalling and reduced cellulose levels has also previously been reported by Ellis et al., (2002). Thus, our finding that *G. hyphopodioides* colonisation results in the upregulation of lignin biosynthesis genes and JA/ET signalling genes is pertinent.

Previous studies have reported higher levels of cell wall lignification in response to colonisation by several non-pathogenic *Magnaporthaceae* species (Huang et al., 2001; Speakman & Lewis, 1978). In the present study, we detected earlier and higher expression of lignin biosynthesis genes in *G. hyphopodioides* colonised tissues compared to *G. tritici* infected tissues. In contrast, local cell wall lignification (as determined by potassium permanganate staining) was more prominent in *G. tritici* infected roots at 5 dpi. However, the downregulation of several COMT genes in response to *G. hyphopodioides* is noteworthy. COMT genes are involved in the synthesis of the S unit of lignin, and downregulation of these genes has a minimal effect on total lignin content (Nguyen et al., 2016). Such changes in lignin composition can drastically alter the outcome of plant-pathogen interactions (Höch et al., 2021; Ma et al., 2018; Quentin et al., 2009). Therefore, despite contrasting results, cell wall lignification could play an important role in *G. hyphopodioides*-mediated take-all control.

In our dataset, 26 TIFY TFs, involved in the cross-talk between JA and other phytohormones (Singh & Mukhopadhyay, 2021) were upregulated in response to *G. hyphopodioides* at 5 dpi. Just three TIFY TFs were significantly upregulated in response to *G. tritici.* In addition, *G. hyphopodioides* colonisation resulted in the upregulation of a greater number of ERF-like genes, known to integrate ET and JA signal transduction pathways (Lorenzo et al., 2003). Phytohormone quantifications using ultra-high-performance liquid chromatography (UHPLC) yielded highly variable results, and we did not detect a significant difference in local JA levels between any treatment. However, high levels of variability may have been due to the transient nature of local JA signalling in plants (Ruan et al., 2019). Thus, unlike in EMR by non-pathogenic *Fusarium* species, *G. hyphopodioides*-induced resistance is potentially mediated, at least to some extent, by the JA/ET signalling pathway. Further investigation is required to determine whether the disruption of CWI mechanisms is directly responsible for the activation of JA/ET mediated defence pathway and the lignin biosynthesis pathway. In addition, future studies should investigate plant and fungal gene expression during *G. tritici* infection of roots already colonised by *G. hyphopodioides*.

In summary, we demonstrate rapid and extensive transcriptional reprogramming in *G. hyphopodioides* colonised wheat roots, characterised by the strong local induction of diverse plant defence mechanisms. We propose that the collective effect of these local defence mechanisms, particularly relating to cell-wall mediated resistance, are responsible for *G. hyphopodioides*-mediated take-all control. Due to the lack of high-quality annotated *G. tritici* and *G. hyphopodioides* genomes, comparative analysis of fungal gene expression during *G. hyphopodioides* colonisation and *G. tritici* infection was not possible in this study. When combined with the RNA-seq dataset presented here, future genome sequencing projects will no doubt facilitate the investigation of novel *G. tritici* pathogenicity factors. In addition, further analysis of non-pathogenic and pathogenic fungi within the diverse *Magnaporthaceae* family may help to address wider questions relating to pathogen organ specificity, conserved fungal root infection strategies and the determinants of fungal pathogenicity.

## Experimental Procedures

### Fungal isolation and culture

*G. hyphopodioides* (taxon id: 1940676) and *G. tritici* (taxon id: 36779) strains were isolated from field soils at Rothamsted Farm using the soil baiting method (McMillan et al., 2011; Osborne et al., 2018). Fungal isolates (see table 1), were maintained on potato dextrose agar (PDA) plates at 21°C in the dark.

**Table 1.**
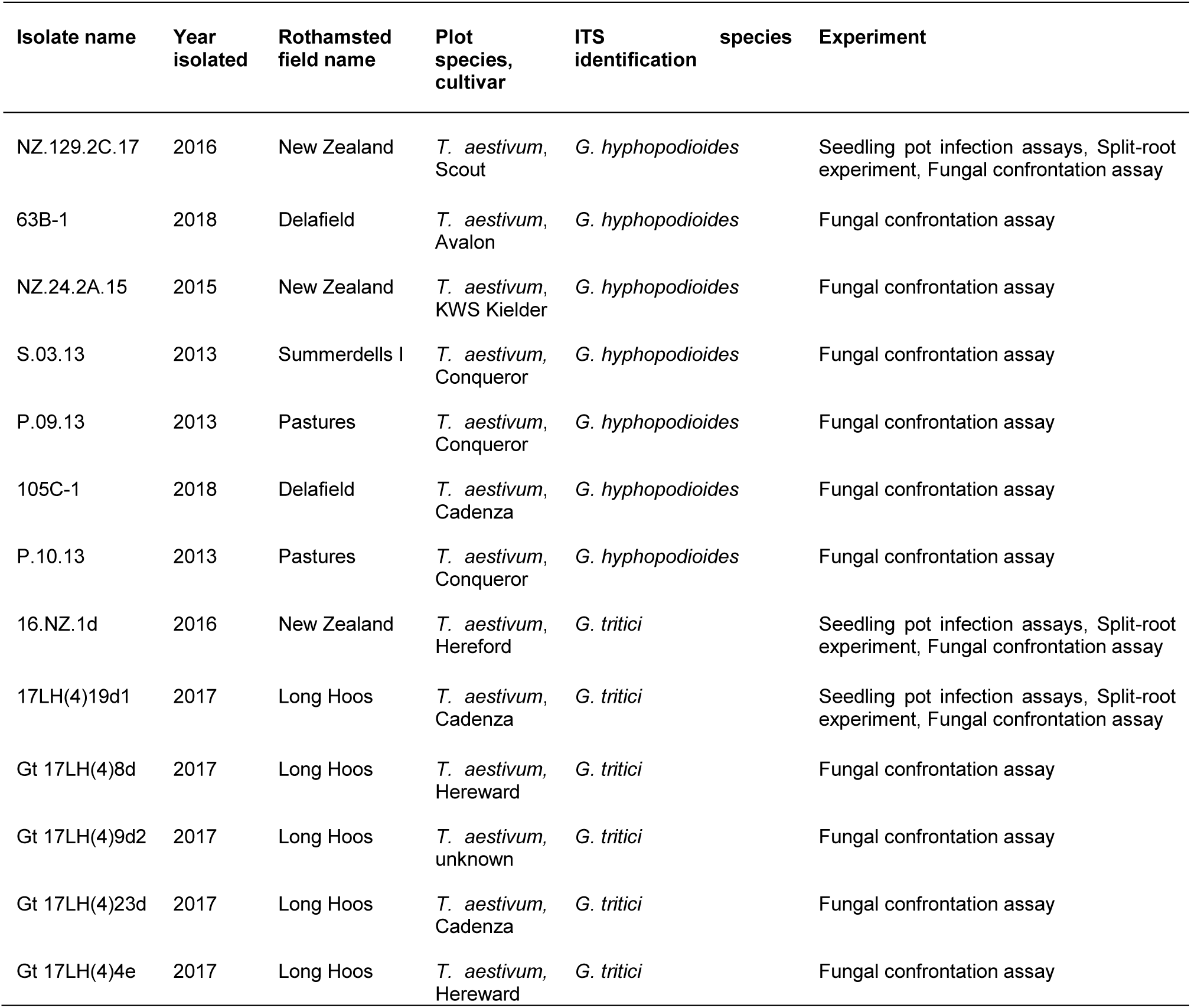
Full list of Rothamsted fungal isolates used in the present study.

### Seedling infection pot assays and disease quantifications

For seedling pot experiments, plastic pots (7.5 cm wide x 11 cm tall) were filled with damp horticultural sand and ten untreated wheat seeds (cv. Hereward) were sown on the surface. Seeds were covered with a thin layer of grit and pots were placed in a controlled environment growth room for two weeks (16 hr day, light intensity 250 µmols, 15 °C day, 10 °C night). *G. tritici* (isolate 16.NZ.1d) and *G. hyphopodioides* (isolate NZ.129.2C.17) inoculum was prepared by placing ten fungal plugs (7 mm diameter) taken from the leading edge of each colony into a 1 L flask containing 400 ml potato dextrose broth (PDB). Flasks were placed in an orbital incubator for 7 days at 25 °C, 120 RPM. Liquid cultures were homogenised by passing through a 2.8 mm sterile sieve. Homogenised cultures were diluted with sterile distilled water in a 2:3 ratio. The first inoculum treatment was added into the pots after two weeks of plant growth. Inoculum (50 ml) was poured directly onto the root system using a funnel inserted into the sand. All seedlings were harvested three weeks after the final inoculum addition to allow take-all disease symptoms to develop (see figure S7A). Five replicates were prepared per treatment, and the experiment was repeated twice.

Visual disease assessments were carried out as previously described (McMillan et al., 2011) and qPCR quantification of *G. tritici* fungal biomass was performed by targeting a 105-bp partial DNA sequence of the translation elongation factor 1-alpha (EF1-α) gene, using primers GtEFF1 (5’-CCCTGCAAGCTCTTCCTCTTAG-3’) and GtEFR1 (5’-GCATGCGAGGTCCCAAAA-3’) with the TaqMan probe (5’-6FAM-ACTGCACAGACCATC-MGB-3’) (Thermo Scientific™, USA) (Keenan et al., 2015).

For split-root experiments, roots from 2-week old wheat seedlings (cv. Chinese Spring) were split across two pots (pot A, pot B) joined at one side. Pots were filled with sand and covered with grit. Roots in pot A received *G. hyphopodioides* liquid inoculum (isolate NZ.129.2C.17), using the method described above. Plants were left to grow for one week before inoculating with *G. tritici* liquid inoculum (isolate 17LH(4)19d1). To investigate whether *G. hyphopodiodes* provides local control against take-all disease, *G. tritici* inoculum was added to *G. hyphopodioides* colonised roots in pot A. To investigate whether *G. hyphopodiodes* provides systemic control against take-all disease, *G. tritici* inoculum was added to uninoculated roots in pot B (see figure S7B). Plants were harvested three weeks later. Five replicates were prepared per treatment, and the experiment was repeated three times.

### Plant growth, inoculation and root sampling for RNA sequencing and bioimaging

A precision inoculation method was developed to enable the investigation of local plant responses to fungal infection (see figure S8). Wheat seeds cv. Chinese Spring were surface sterilised with 5% (v/v) sodium hypochlorite for five minutes and pre-germinated in a controlled environment growth chamber cabinet (20 °C day, 16 °C night, 16 hr light cycle) for two days. Three pre-germinated seeds were transplanted onto a square petri dish plate (12 cm x 12 cm) containing 1.5 % (w/v) water agar. Five replicates were prepared for each treatment. Plates were placed vertically in the growth cabinet. After four days, one root from each plant was inoculated with a fungal plug (4 cm x 0.5 cm) cut from the leading edge of a 2-week old fungal colony growing on 1.5 % water agar. Inoculated roots were sampled daily from 2-6 days post inoculation (dpi). Briefly, two 1 cm root samples were harvested from the inoculated area on each root and snap frozen in liquid nitrogen for RNA extraction. To determine the stage of fungal colonisation in these harvested samples, 2 x 0.5 cm root pieces were sampled from the areas directly above and below each sample. Root pieces were stored in 50% ethanol for subsequent assessment by confocal microscopy.

### Fluorescent staining and confocal microscopy analyses

To assess colonisation in whole root pieces, samples were cleared in 10% w/v potassium hydroxide for 5 minutes at 70 °C, before staining with Propidium Iodide (PI) (10 µg/ml) and Wheat Germ Agglutinin, Alexa Fluor™ 488 Conjugate (WGA) (10 µg/ml). To visualise vascular infection by *G. tritici*, transversal root sections were cut by hand using a fine edged razor blade under a dissecting microscope. Confocal microscopy was performed using a ZEISS 780 Confocal Laser Scanning Microscope (ZEISS, Germany). WGA fluorescence was excited at 495 nm and detected at 519 nm. PI fluorescence was excited at 535 nm and detected at 617 nm.

### RNA extraction

Following confocal assessment (see above), root pieces (1 cm each) at the same stage of fungal colonisation were pooled together to create a single sample for RNA extraction. Total RNA was extracted from frozen root material using the E.Z.N.A.® Plant RNA Kit with the associated RNase-free DNase I Set (Omega-Biotek, USA), following the standard protocol provided. RNA quality was assessed based on the RNA Integrity Number (RIN), measured using the Bioanalyser 2100 with the corresponding RNA 6000 Nano Kit (Agilent, USA), as per manufacturer instructions.

### Library preparation and sequencing

mRNA library preparation was carried out by Novogene (China) using the Novogene RNA Library Prep Set (PT042) for polyA enrichment. Libraries were sequenced by Illumina NovaSeq to generate 150 bp paired-end reads, with a target of 40 million paired-end reads per sample.

### Transcriptome annotation and analysis

Quality control of reads was performed using MultiQC (https://multiqc.info/). Sequence trimming of recognised adaptors was performed using Trimmomatic where appropriate (Bolger et al., 2014). Reads were mapped to the Chinese Spring (IWGSC RefSeq v2.1) (Zhu et al., 2021) using HiSat2 (Kim et al., 2019). To ensure that fungal biomass was consistent among replicates of the same treatment, reads were also mapped to the *G. tritici* genome (Okagaki et al., 2015). Three samples were identified as outliers based on standardised residuals of the percentage of reads mapped to *G. tritici.* Outliers were subsequently excluded from further analyses (table S10). All treatments contained at least four biological replicates, with the majority containing five biological replicates. Reads were not aligned to *G. hyphopodioides* due to the lack of a high-quality genome. Count determination was performed using FeatureCounts (Liao et al., 2014) on the R Bioconductor platform (https://bioconductor.org/).

Library normalisation and differential expression (DE) calling was carried out using the Bioconductor package DESeq2 (Love et al., 2014) in R studio. Gene expression levels were compared between *G. tritici* infected and *G. hyphopodioides* colonised samples and the uninoculated control samples for each time point individually. DE genes were identified by applying a log2 fold change filter of ≥ 1 or ≤ −1. The DESeq2 implementation of Benjamini-Hochberg (Benjamini & Hochberg, 1995) was used to control for multiple testing (q<0.05). Gene Ontology (GO) enrichment analysis was performed for significantly up- and down- regulated wheat genes separately via http://www.geneontology.org, using the Panther classification system.

### Statistical analyses

Statistical analyses were done using Genstat 20th Edition (VSN International Ltd, Hemel Hempstead, UK). Percentage disease data were analysed using a Generalised Linear Regression Model (GLM) with a binomial distribution and LOGIT link function. Analyses were adjusted for over-dispersion and treatment effects tested using deviance ratios (F-statistics) when the residual mean deviance was greater than 1. Data were back-transformed from the LOGIT scale (using the equation exp(x)/(1+exp(x))) for graphical presentation. For continuous outcome variables such as plant biomass, data were analysed by Analysis of Variance (ANOVA). Tukey’s multiple comparisons test was carried out when more than one interaction was of interest.

## Acknowledgements

TC was supported by the Biotechnology and Biological Sciences Research Council (BBSRC) funded University of Nottingham Doctoral Training Programme (BB/M008770/1). JPG, DS and KHK were supported by the BBSRC Institute Strategic Programme (ISP) Grant, Designing Future Wheat (BBS/E/C/000I0250). In addition, DS is supported by the BBSRC ISP Grant (BB/CCG2280/1) and KHK by the BBSRC ISP Grant, Delivering Sustainable Wheat (BB/X011003/1 and BBS/E/RH/230001B). GC is supported by the defra funded Wheat Genetic Improvement Network, WGIN (CH0109). VA is supported by the BBSRC funded South West Biosciences Doctoral Training Partnership (BB/T008741/1).

We thank Smita Kurup and Hannah Walpole (Plant Sciences for the Bioeconomy, Rothamsted Research) for advice and assistance with fluorescent staining and confocal microscopy of wheat roots; Jess Hammond (Protecting Crops and the Environment, Rothamsted Research) for help with fungal isolations and plant care; Matthew Dickinson (University of Nottingham) for his feedback on the project.

## Data availability statement

The data that support the findings of this study are openly available in the NCBI Gene Expression Omnibus (GEO) at https://www.ncbi.nlm.nih.gov/geo/, reference number GSE242417.

