## Supplementary figures and methods for "A fungal endophyte induces local cell-wall mediated resistance in wheat roots against take-all disease"

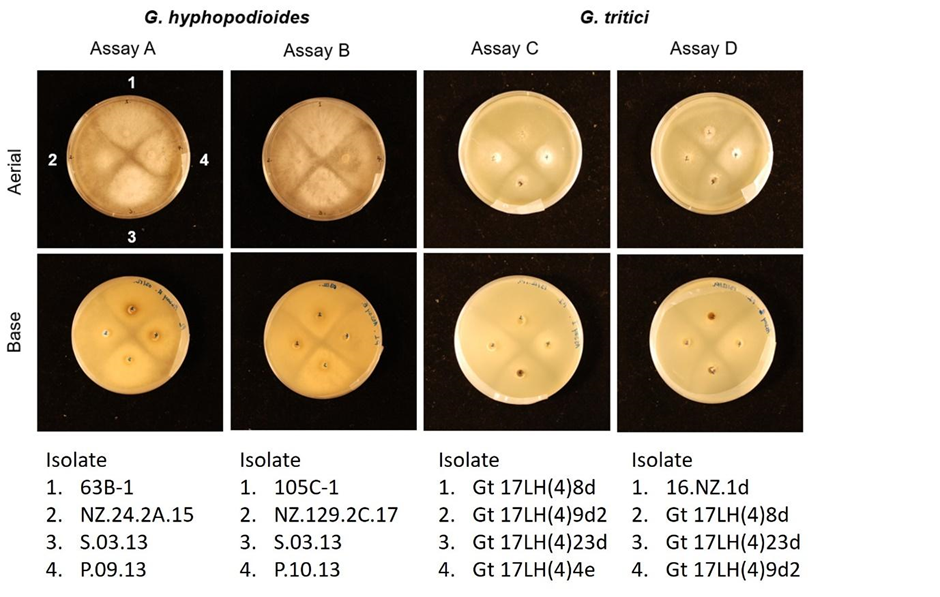

**Figure S1.** *G. hyphopodioides* and *G. tritici* intra-species confrontation assays on potato dextrose agar (PDA). Imaged at 11 days of fungal growth. The locations of fungal isolates have been indicated by position numbers 1-4.

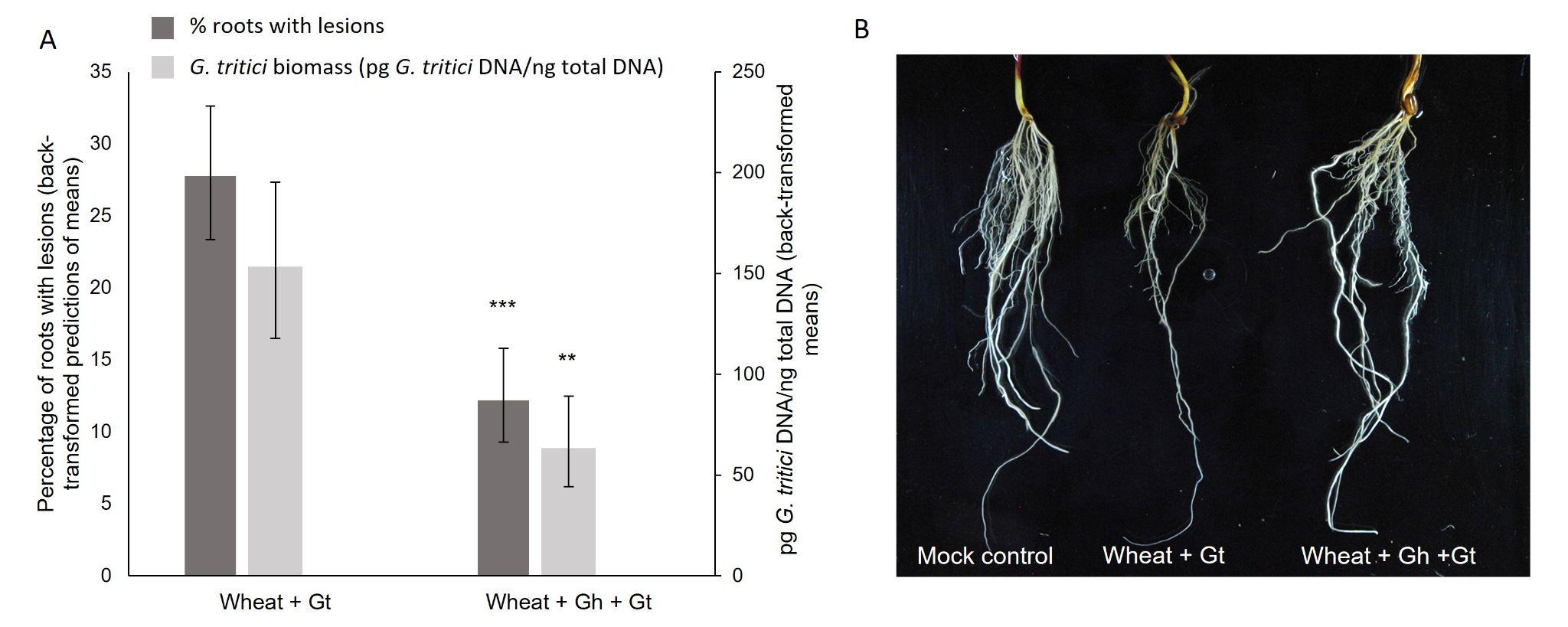

**Figure S2.** **Additional wheat (cv. Chinese Spring) co-inoculation experiment with *G. tritici* (17LH(4)19d1) and *G. hyphopodioides* (NZ.129.2C.17), whereby *G. hyphopodioides* was added to pots one-week prior to *G. tritici.*** A. The percentage of wheat roots with take-all lesions and amount of *G. tritici* fungal biomass (pg *G. tritici* DNA/ ng total DNA) with and without *G. hyphopodioides* pre-treatment. The data revealed a clear reduction in both take-all disease levels (GLM: d.f. 1, 21, F=29.24, p<0.001) and *G. tritici* fungal biomass (ANOVA: d.f. 1, 8, F=15.54, p=0.004) in plants pre-treated with *G. hyphopodioides;*  B. Representative images of mock inoculated roots, roots infected with *G. tritici* only, and roots infected with *G. tritici* following *G. hyphopodioides* pre-treatment.

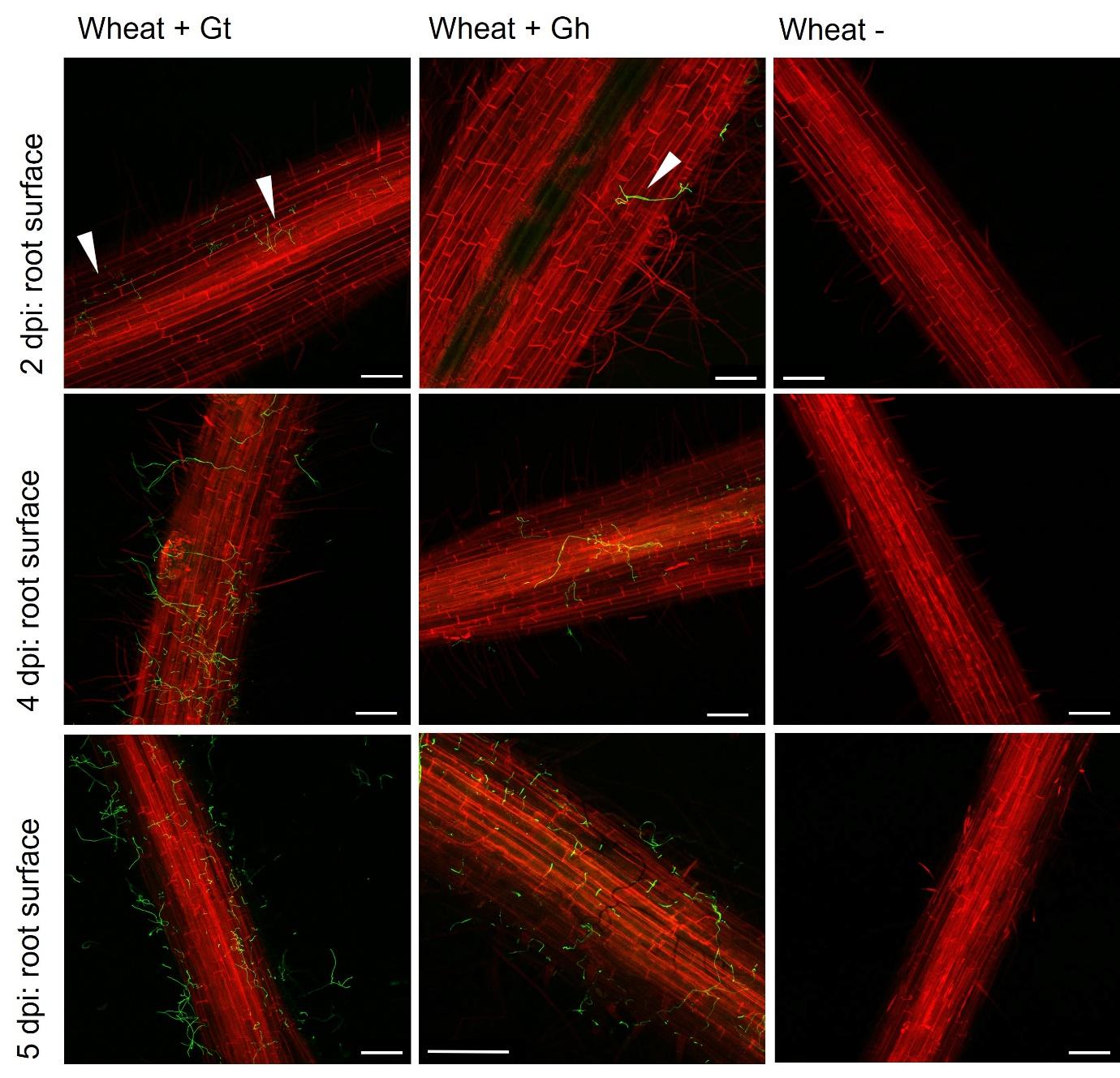

**Figure S3. Fluorescence images obtained by confocal microscopy showing either *G. tritici* (17LH(4)19d1) or *G. hyphopodioides* (NZ.129.2C.17) hyphae on the surface of wheat roots (cv. Chinese Spring) at key time points selected for RNA sequencing.** White arrows indicate fungal hyphae. Scale bars represent 200 µm.

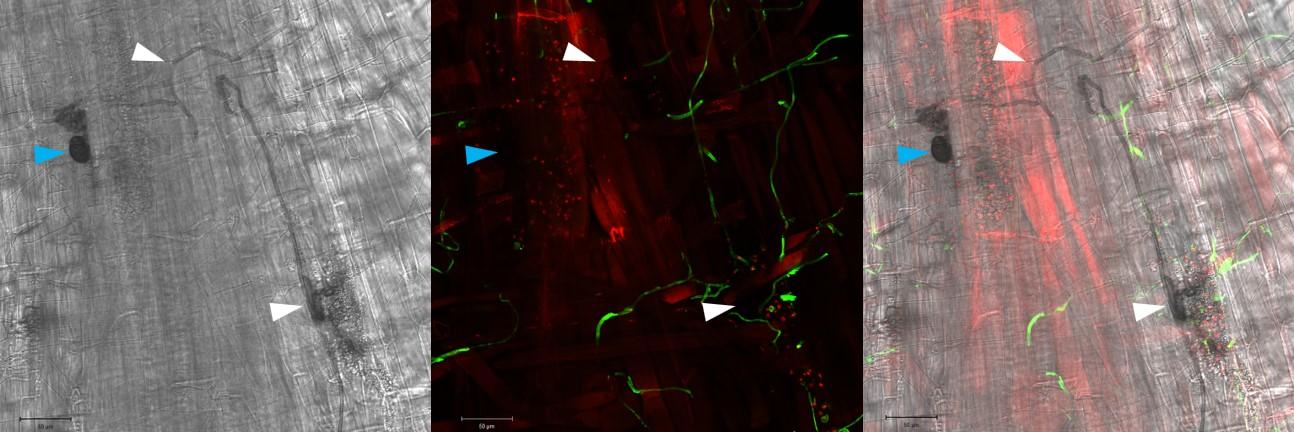

**Figure S4. Fluorescence and transmitted light image overlays showing *G. hyphopodioides* SEVs and macrohyphae with and without WGA staining at 5 dpi.** Blue arrows indicate unstained SEVs. White arrows indicate unstained macrohyphae. Scale bars represent 50 µm.

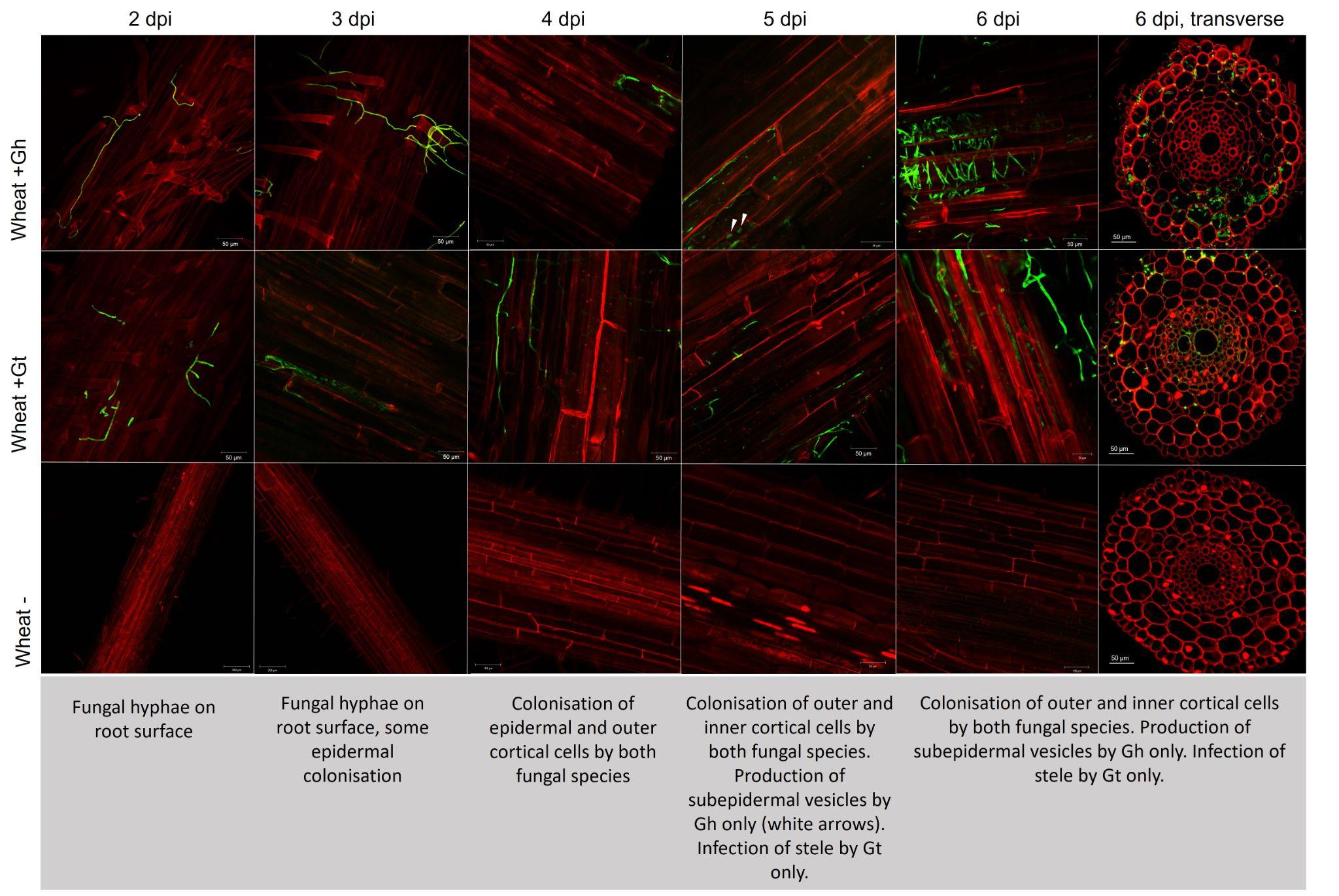

**Figure S5. Representative confocal micrographs of wheat roots (cv. Chinese Spring) infected with either *G. tritici* or *G. hyphopodioides* at 2, 4, 5 and 6 dpi.** White arrows indicate newly formed SEVs.

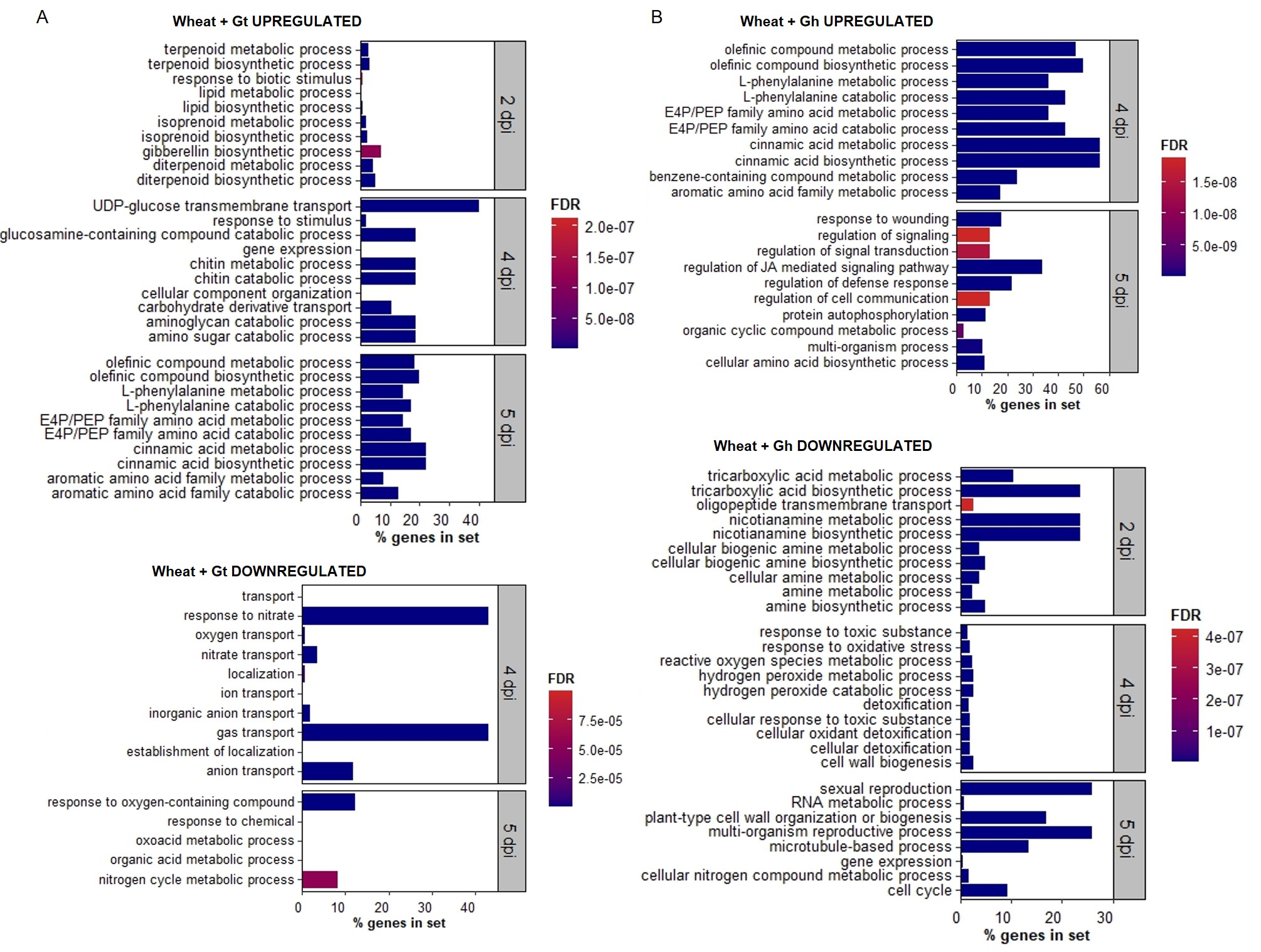

**Figure S6. Wheat transcriptional changes during the infection progression of *G. hyphopodioides* compared to *G. tritici.*** A. Top ten newly enriched biological process GO terms in *G. tritici* infected root samples compared to uninoculated controls; B. Top ten newly enriched biological process GO terms in *G. hyphopodioides* colonised root samples compared to uninoculated controls. To identify the top 10 newly enriched terms, biological process GO terms were compared between each time point and the top ten newly enriched terms were identified. Percentage of genes in a set represents the percentage of DE genes out of the total number of known genes in that category, according to the Panther classification system.

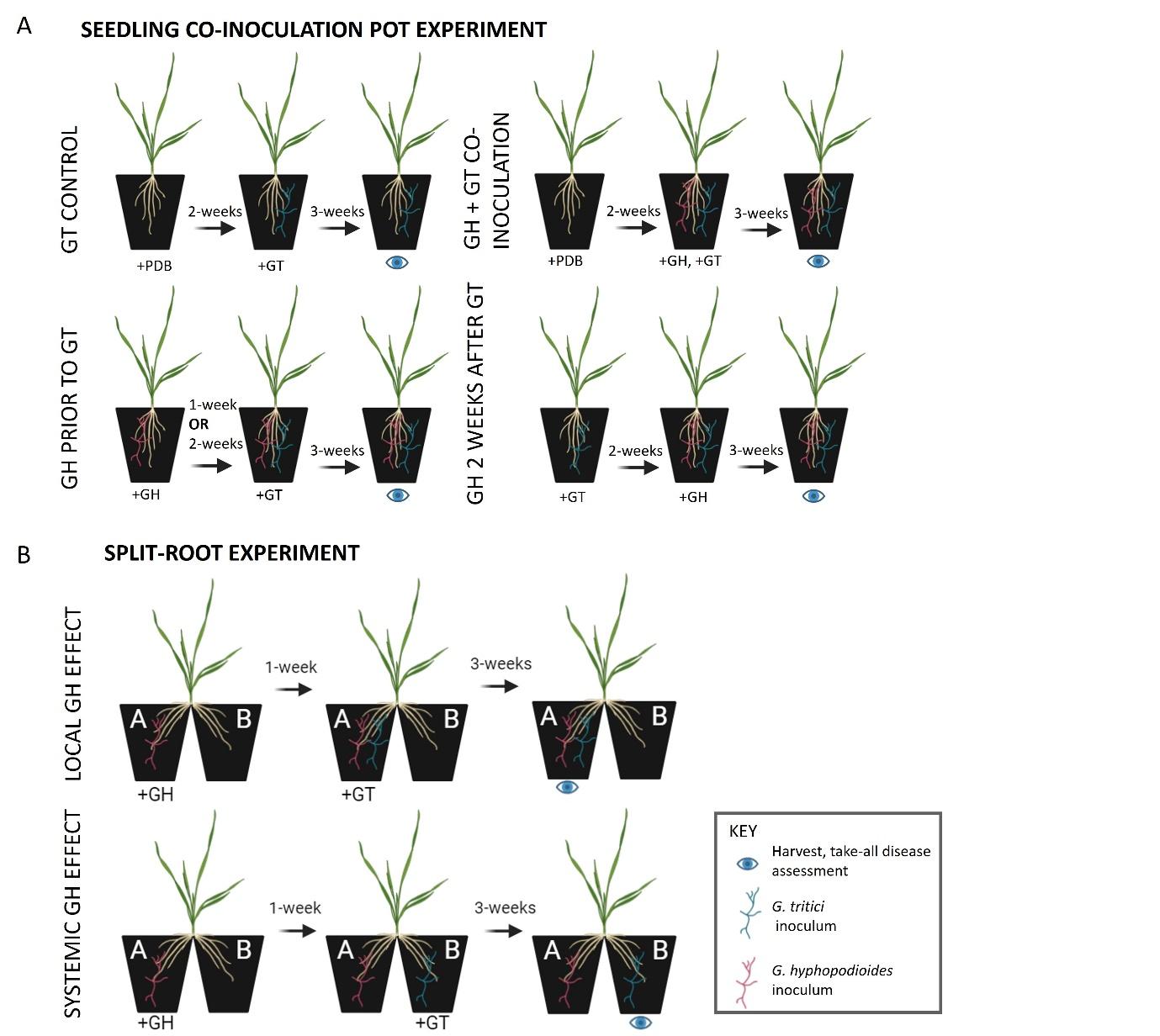

**Figure S7. Schematic of pot inoculation experiments**. A. Wheat seedling co-inoculation assays with *G. hyphopodioides* (Gt) and *G. tritici* (Gh)*. Gh* inoculum was added either 2-weeks prior, 1-week prior, at the same time as, or 2-weeks after *Gt* inoculum; B. Split root experiments. To investigate the local effect of *Gh* colonisation on take-all disease levels, *Gt* inoculum was added to roots in pot A, which had been pre-treated with *Gh* 1-week prior. To investigate the systemic effect of *Gh* colonisation on take-all disease levels, *Gt* was added to roots in pot B, which had not been pre-treated with *Gh*. All plants were harvested for take-all disease assessments 3-weeks after the final inoculum addition. Created in BioRender.com.

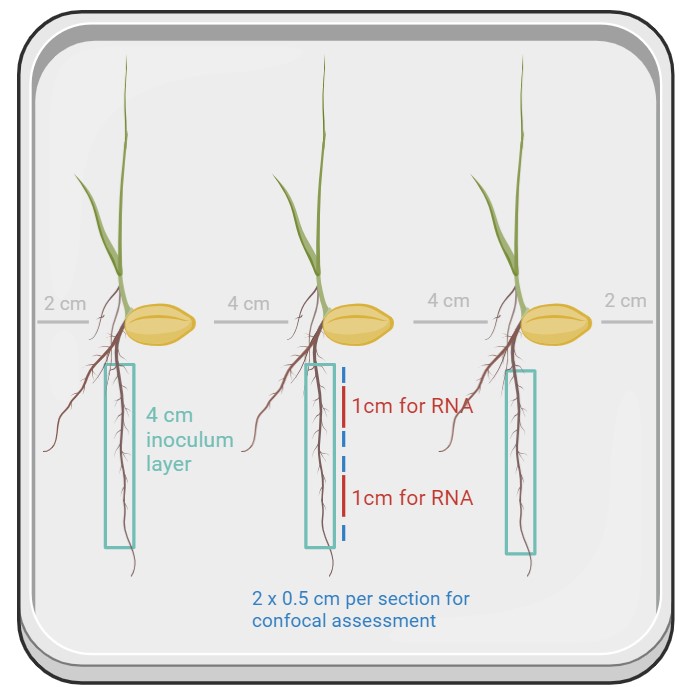

**Figure S8. Schematic showing the precision inoculation method used to investigate the wheat response to *G. hyphopodioides* colonisation or *G. tritici* infection.** Water agar plates were sown with 3x pre-germinated wheat seedlings (cv. Chinese Spring). One root per plant was inoculated with a 4 cm inoculum plug from either a *G. tritici* or a *G. hyphopodioides* fungal colony. To investigate the local plant response to fungal infection, two 1 cm root samples were harvested from the inoculated area on each root and snap frozen in liquid nitrogen for RNA extraction. To determine the stage of fungal colonisation in these harvested samples, 2 x 0.5 cm root pieces were sampled from the areas directly above and below each sample. Created in BioRender.com.

### Supplementary Methods

**Transmission electron microscopy of G. hyphopodioides subepidermal vesicles (SEVs)**

*G. hyphopodioides* colonised root tissues were harvested from a seedling infection pot assay at 5 weeks post inoculation. Roots were screened under a Leica M205 stereomicroscope (Leica Microsystems) for the presence of mature SEVs, as indicated by a highly melanised cell wall. Root pieces containing mature SEVs were excised and prepared for transmission electron microscopy (TEM) analysis by high-pressure freezing (Leica HPM100), followed by freeze substitution with ethanol (Leica EM AFS2) and infiltration with LR White resin (Agar Scientific). Ultrathin sections (90 nm thickness) were cut using a UC7 Ultramicrotome (Leica Microsystems). Due to the large size of *G. hyphopodioides* SEVs, a subset of semi-thin sections were stained and analysed by light microscopy to allow visualisation of entire SEVs (see below for staining conditions).

The remaining ultrathin sections were collected onto copper grids and stained with 2.5% uranyl acetate and Reynolds lead citrate (Reynolds, 1963). Ultrathin sections were imaged using a JEOL-2100 Plus Transmission Electron Microscope (JEOL, Japan) equipped with a Gatan OneView IS Camera (Gatan, USA). ImageJ software (<https://imagej.net/>) was used to measure the cell wall diameter of *G. hyphopodioides* SEVs and infection hyphae. The average of ten measurements, taken at random across the cell wall, was calculated for three SEVs and three hyphae in each plant sample. Measurements were repeated across three different plant samples.

**Light Microscopy**

Root samples were placed in fixative at 4 °C overnight (4% paraformaldehyde, 2.5% glutaraldehyde in 0.05M phosphate buffer, pH 7.2), before dehydration in a graded ethanol series. Roots were infiltrated and embedded in glycol methacrylate resin using the Agar GMA HEMA kit (Agar Scientific, UK). Embedded samples were sectioned to 4 µm thickness. To visualise plant lignin, sections were stained with 1% w/v potassium permanganate for 10 minutes. To visualise fungal structures, sections were stained with 1% w/v aniline blue in lactophenol for 30 seconds. Semi-thin LR White resin sections containing *G. hyphopodioides* SEVs (see above) were stained with 1% w/v toluidine blue for 1 minute. All stained sections were mounted onto slides with DPX mounting fluid (Sigma Aldrich®). Sections were imaged using a ZEISS Axio Imager light microscope (ZEISS, Germany).

**Validation of gene expression by RT-qPCR**

The transcript abundance of selected DEGs was assessed in 5 dpi samples by RT-qPCR. cDNA was synthesised using the SuperScript™ IV Reverse Transcriptase Kit with Oligo (dT) primers (ThermoFisher Scientific, USA), according to the manufacturer’s instructions. RT-qPCR was carried out using the PowerTrack™ SYBR Green Master Mix (ThermoFisher Scientific, USA) on the QuantStudio™ 6 Pro Real-Time PCR System (Applied Biosystems™, USA). Relative gene expression levels were calculated using the 2–∆∆CT (Livak) method (Livak & Schmittgen, 2001), following normalisation to the cell division control 48 (CDC48) housekeeping gene (Lee et al., 2014). Primer sequences used in qPCR assays can be found in table S11.

**Phytohormone quantification**

To generate sufficient root material (30 mg dry weight) for the quantification of local JA and SA levels, an experimental repeat was carried out using the precision inoculation method. Roots were sampled at 5 dpi by harvesting the 4 cm inoculated root area from each plant. Samples were pooled and lyophilised for 24 hours in a LyoDry Compact Benchtop Freeze Dryer (Mechatech systems, UK). Samples were ground to a fine powder in a Geno/Grinder® (SPEX Europe, UK). Hormones were separated by UHPLC with a reverse Accucore C18 column (2.6 µm, 100 mm length; Thermo Scientific™) and an acetonitrile gradient containing 0.05% acetic acid. The hormones were analysed with a Q-Exactive mass spectrometer (Orbitrap detector; Thermo Scientific™). The concentrations of hormones in the extracts were determined using embedded calibration curves and the Xcalibur 4.0 and TraceFinder 4.1 SP1 programs. Three biological replicates per treatment were analysed.

**Lignin quantification**

In order to ascertain the relative level of lignification in root tissue samples embedded in resin, potassium permanganate staining was used. For the analysis of potassium permanganate-stained root tissues, image scale was set using the ‘set scale’ function in Fiji (ImageJ) and cropped to area of interest to remove excess background. Images were then parsed using the threshold colour function in the HSB colour space with the parameters of: Hue 0-230, Saturation 0-255, Brightness 0-210, to remove lightly stained fungal hyphae and background from downstream analysis. Using the selected area, the total area of the cell wall was recorded using the measure function. Subsequently, a second measurement of the image was taken to determine the proportion of the cell walls stained darker, indicating a greater lignin presence, using the thresholding parameters of: Hue 0-230, Saturation 0-255, Brightness 0-160. The percentage of the total area stained dark was calculated.
